# Sex-specific developmental phenotypes and their response to neonatal *Dyrk1a* reduction in the Ts65Dn Down syndrome mouse model

**DOI:** 10.64898/2026.07.15.738703

**Authors:** Alyssa Duerst, Laura Hawley, Linnea Johnson, Zaina Obeid, Laura Snellenberger, Drew Folz, Charles R. Goodlett, Randall J. Roper

**Affiliations:** Department of Biology, Indiana University Indianapolis, 723 W Michigan Street, SL306, Indianapolis, IN 46202, USA; Department of Psychology, Indiana University Indianapolis, 402 N Blackford St, LD124, Indianapolis, IN 46202, USA; Stark Neuroscience Research Institute, Indiana University School of Medicine, Indianapolis, Indiana, USA

**Author notes:** Corresponding Author Randall J. Roper. These authors contributed equally to this work (Alyssa Duerst, Laura Hawley).

**Keywords:** Trisomy 21, development, *Dyrk1a*, sex differences, behavior

## Abstract

Children with Down syndrome (DS) experience delays in cognitive, physical, and motor development. Overexpression of *Dual-specificity tyrosine phosphorylation-regulated kinase-1A* (*DYRK1A*), a gene on human chromosome 21 (Hsa21) and triplicated in individuals with Trisomy 21, contributes to neurodevelopmental delays associated with DS, and is a candidate for therapies to improve neurodevelopmental phenotypes. Male and female Ts65Dn DS model pups are trisomic for ∼100 Hsa21 orthologs including *Dyrk1a,* and both sexes show significant DYRK1A overexpression on postnatal day 6 (P6) in the hippocampus, cerebral cortex, and cerebellum. This study tested the hypothesis that normalization of *Dyrk1a* copy number in Ts65Dn pups prior to P6 would diminish physical and behavioral developmental outcomes in Ts65Dn mice, thus providing a standard of comparison for success of interventions targeting *Dyrk1a.* At P3-P21, Ts65Dn compared to euploid pups showed sex-specific deficits in physical, motor, and behavioral development. Male Ts,*Dyrk1a*^+/+/Dox-Cre^ mice showed improved emergence to running on P19, and both sexes of Ts,*Dyrk1a*^+/+/Dox-Cre^ mice exhibited reduced isolation-induced ultrasonic vocalizations during the second postnatal week. *Dyrk1a* normalization in Ts65Dn pups did not improve all abnormal phenotypes, perhaps because of developmental dysregulation between *Dyrk1a* RNA and DYRK1A protein levels, involvement of other trisomic genes, or improvements in only adult mice.

## INTRODUCTION

The partial or full triplication of human chromosome 21 (Hsa21) results in Down syndrome (DS), and developmental and cognitive deficits in varying degrees are prominent among the many phenotypes expressed [1]. Dosage imbalance of genes on Hsa21 causes growth deficits, behavioral abnormalities, and altered structural brain phenotypes associated with interrupted proliferation and differentiation of cells and disrupted neuronal network connectivity that hamper information processing in the brain [2,3]. Developmental delays are a hallmark of DS. Around the age of 2, the disparity between affected individuals and typically developing peers becomes more apparent [4–7]. Neurodevelopmental abnormalities in toddlers and children with DS are associated with communication, motor, and learning delays [6,8]. Typical DS phenotypes, such as hypotonia and altered sensorimotor development, present as delays in motor skills such as sitting, crawling, and walking in affected individuals [9–11].

Sexual dimorphisms in deficits have been observed in individuals with Ts21. Developmental milestones are generally achieved earlier in females than males with DS, though in some motor milestones, males with DS begin to complete the task before females (using stairs, standing, and scooting, creeping or crawling) [11]. Girls with DS are more likely to show higher intelligence, adaptive function, and developmental age compared to boys [12,13]. During adolescence, females exhibit fewer behavioral problems than males [14]. In adults, women show higher cognitive abilities compared to men and the frequency of profound intellectual disability was twice as high in men as women [15]. Understanding the origins and developmental trajectories of phenotypic differences between males and females with DS is important for developing potential interventions to improve sex-specific deficits.

DS mouse models have been utilized to understand structural and behavioral changes caused by dosage imbalance of genes orthologous to those on Hsa21. The Ts(17^16^)65Dn (Ts65Dn) DS mouse model contains a third copy of about half of Hsa21 orthologs on a freely segregating, segmental chromosome made up of the distal end of mouse chromosome (Mmu) 16 attached to a Mmu17 centromere [6,16,17], though it lacks the complete complement of Hsa21 orthologs and is trisomic for ∼35 protein coding genes not homologous to Hsa21 [18,19]. This mouse model has been used in testing intervention therapies designed to improve DS phenotypes [17,20–23]. Ts65Dn as compared to littermate control mice present with early developmental delays that model individuals with DS, including lower body weight and slowed weight gain during development [24–32]. Structural deficits, such as fewer cerebellar proliferating granule cell precursors with elongated cell cycles and hypomorphic brain composition, contribute to the cognitive impairments that are indicative of Ts21 [3,17,20,23,33–35].

Many early studies using DS mouse models only analyzed male Ts65Dn mice because female mice were reserved for colony maintenance, given that these male mice are largely infertile. During early postnatal development, male Ts65Dn mice (of the original Ts65Dn 001924 line) showed reduced weight gain and delays in sensorimotor development, but no delays in eye opening or incisor eruption [25]. Ultrasonic vocalizations (USVs) elicited by separation from the dam were delayed in Ts65Dn pups; peak rate of USV calls emitted by euploid mice occurred on postnatal (P) day 6 and declined thereafter, whereas for trisomic mice, the peak rate occurred on P9 before declining. Homing behavior was abnormal in trisomic as compared to euploid mice, and open field activity was elevated both on P21 and in adult mice. A recent study of early postnatal behavior development in male and female Ts65Dn mice (using the 005252 line) found no significant differences between trisomic male and female pups in several typical DS phenotypes, such as reduced weight gain, delayed sensorimotor task acquisition, and cliff aversion. However, significant differences were present between the sexes of trisomic pups in achieving growth milestones, with male mice exhibiting more delayed development than female mice [32].

*DYRK1A*, one of the genes triplicated in Ts21 and Ts65Dn mice, is a serine-threonine kinase that is implicated in multiple pathways of neurogenesis and neurodevelopment, playing a critical role in transcriptional regulation of genes responsible for cell cycle progression, neuronal proliferation, and cellular differentiation [36–44]. *Dyrk1a* is highly conserved between species and is critical for neurodevelopment. During typical murine neurodevelopment, DYRK1A is produced in a regulated pattern of expression and localization [45]. In situ hybridization of *Dyrk1a* mRNA in brain tissues of typically developing embryonic mice found spatiotemporal patterns of expression coinciding with active neuron maturation [46]. Between P7-P14, *Dyrk1a* overexpressing mice (the Tg*Dyrk1A* model) showed delays in rostro-caudal maturation of the motor system, which may have led to motor deficit phenotypes including walking, pivoting, and homing [47]. Haploinsufficiency of *DYRK1A* in humans and mice leads to phenotypes associated with DYRK1A syndrome [48,49], providing additional evidence of the importance of DYRK1A levels in regulating developmental processes.

Conventionally, it has been assumed that an increased gene copy number will cause a proportionally higher level of gene expression, yet evidence has shown that this is not always true. In the first three weeks after birth, DYRK1A protein expression was significantly dysregulated in Ts65Dn compared to euploid littermate control pups, with fluctuations that depended on the age, sex, and brain region of the subject [50]. Ts65Dn mice showed significant overexpression of DYRK1A on P6 in the cerebellum (CB), hippocampus (HIP), and cerebral cortex (CTX) of both male and female pups, an age of significant neuronal (granule cell) and glial proliferation [51]. In males, this overexpression continued through P15, 18, 21, and 24; in contrast, DYRK1A was not significantly overexpressed in female Ts65Dn mice from P15-24 [50]. Given the known detrimental effects of trisomic *Dyrk1a* expression on neurodevelopment, these sex- and age-specific patterns of *Dyrk1a* expression in Ts65Dn mice are relevant to potential treatments targeting DYRK1A activity. Understanding the developmental landscape of *Dyrk1a* expression is necessary to guide appropriate treatment intervention times while avoiding detrimental effects of over-inhibition of DYRK1A and the potential consequences predicted by DYRK1A haploinsufficiency [52]. Preclinical studies of early postnatal pharmacological interventions in DS mouse models provide an important means to assess the efficacy of DYRK1A inhibition to realign growth and structural trajectories before and during emergence of neurobehavioral abnormalities [52–54].

Crossing Ts65Dn females with *Dyrk1a* haploinsufficient males produced some offspring with normalized *Dyrk1a* gene copy number from conception in otherwise trisomic mice. These male Ts65Dn,*Dyrk1a*^+/+/-^ mice showed improved spatial and working memory and partially restored reference memory, improved contextual but not cued fear conditioning, and increased hippocampal long-term potentiation as compared to Ts65Dn,*Dyrk1a*^+/+/+^ mice [55]. Normalization of *Dyrk1a* from conception in Ts65Dn mice did not ameliorate deficits in body weight, motor function, motor coordination, general activity, or anxiety found in Ts65Dn mice. Gait was improved, with stride length and stride number (but not stride width) being normalized in male Ts65Dn,*Dyrk1a*^+/+/-^ as compared to Ts65Dn mice with three copies of *Dyrk1a* [21]. The improvement of some but not all cognitive and behavioral phenotypes resulting from normalizing *Dyrk1a* copy number from conception in Ts65Dn mice gives impetus to develop strategies to ameliorate deficits caused by the developmental effects of triplicated *Dyrk1a*.

To understand the contribution of three copies of *DYRK1A* to early developmental phenotypes of DS and, by inference, how a DYRK1A inhibitor given during development could potentially affect those phenotypes, a DS mouse model with a temporally specific *Dyrk1a* functional reduction was produced. It was hypothesized that inducing a functional reduction of *Dyrk1a* gene copy number in Ts65Dn and normalizing *Dyrk1a* copy number prior to a point of known DYRK1A overexpression would improve developmental phenotypes in DS mice in a sex-specific manner [52]. The temporally specific normalization of *Dyrk1a* would provide a genetic basis for potential use of DYRK1A inhibitors during development to mitigate emerging behavioral deficits and improve developmental trajectories in individuals with DS.

The genetic approach used in this experiment provided the basis to test three a priori hypotheses regarding typical postnatal developmental benchmarks: 1) trisomic mice, as compared to euploid mice of the same sex, will display deficits in growth along with developmental delays in achieving physical and sensorimotor milestones, in the ontogeny of isolation-induced ultrasonic vocalizations, in maturation of motor function, and in emergence of locomotor activity, and will have age-dependent overexpression of DYRK1A in select brain tissues; 2) an early postnatal functional reduction of *Dyrk1a* in otherwise trisomic mice will diminish the molecular, developmental, and behavioral abnormalities as compared to trisomic mice in a sex-specific manner; and 3) a reduction of *Dyrk1a* in otherwise euploid mice (haploinsufficiency) will cause abnormalities in developmental phenotypes as compared to euploid mice, consistent with DYRK1A syndrome [49,56]. Results from this early postnatal genetic intervention therapy on molecular and behavioral phenotypes would help determine if measurable differences in growth and sensorimotor development compared to neurotypical littermate controls could be realized by therapeutic DYRK1A normalization during this window of development.

## RESULTS

### Postnatal body weight

Throughout the postnatal period (P3-21), Ts,*Dyrk1a*^+/+/+^ pups of both sexes weighed significantly less than same-sex Eu,*Dyrk1a*^+*/*+^ littermates (Fig. 1). Additionally, female Eu,*Dyrk1a*^+*/*Dox-Cre^ pups weighed significantly less than their female Eu,*Dyrk1a*^+*/*+^ littermates beginning at P10 and continuing to P20, whereas reductions in male Eu,*Dyrk1a*^+*/*Dox-Cre^ mice relative to Eu,*Dyrk1a*^+*/*+^ males reached significance only on P20 and P21. Functional reduction of *Dyrk1a* in Ts,*Dyrk1a*^+/+/Dox-Cre^ mice failed to improve the postnatal weight deficits in Ts65Dn mice. Daily weight gain (shown in Supplemental Fig. 5 as percentage increase from previous day) was greatest in all groups over the first postnatal week, but Ts,*Dyrk1a*^+/+/+^ mice of both sexes had significantly less daily weight increase than same-sex Eu,*Dyrk1a*^+/+^ littermates during that week. In addition, Ts,*Dyrk1a*^+/+/+^ mice of both sexes had significantly less daily weight gain than same-sex Eu,*Dyrk1a*^+/+^ littermates between P17-P21.

**Figure 1:**
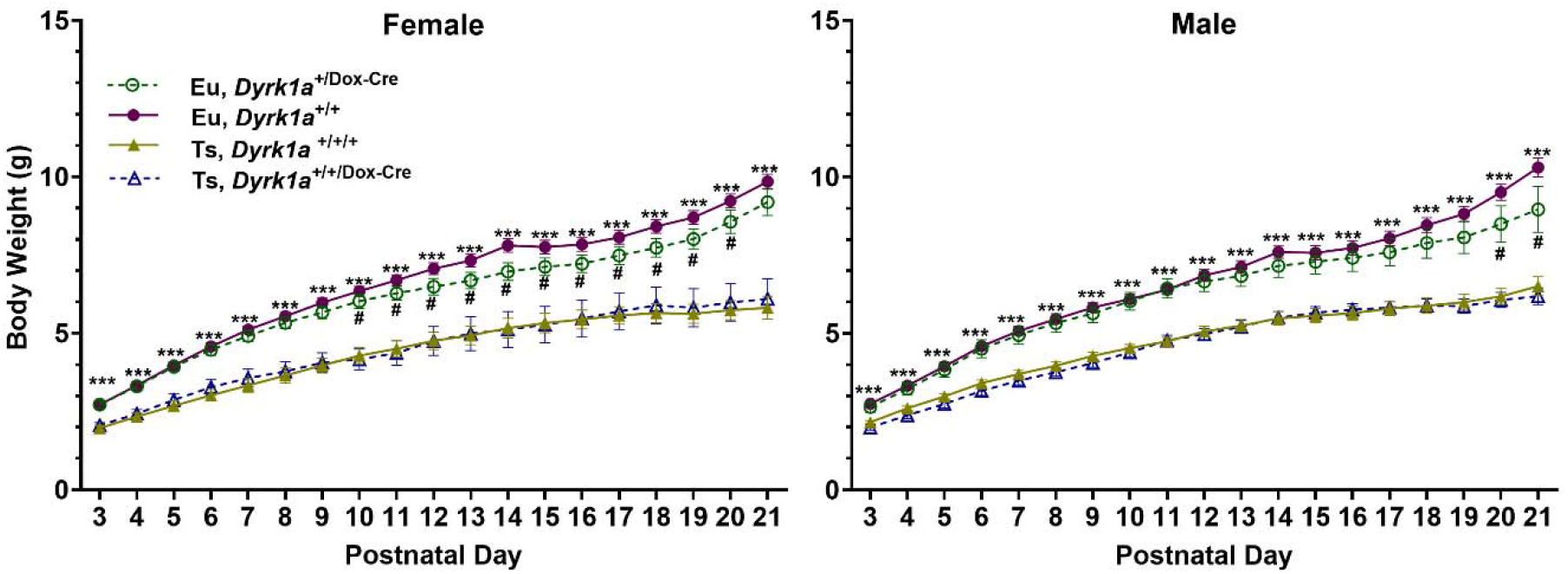
Daily Body Weight of Female and Male Pups. In both sexes, trisomic mice (Ts,*Dyrk1a*^+/+/+^) weighed significantly less than euploid mice (Eu,*Dyrk1a*^+/+^) throughout the postnatal period; functional reduction of *Dyrk1a* (Ts,*Dyrk1a*^+/+/Dox-Cre^) did not improve the postnatal growth deficits in Ts65Dn mice. Female euploid mice with one copy of *Dyrk1a* (Eu,*Dyrk1a*^+/Dox-Cre^)weighed significantly less than female Eu,*Dyrk1a*^+/+^ mice beginning at P10. Male Eu,*Dyrk1a*^+/Dox-Cre^ mice weighed significantly less than Eu,*Dyrk1a*^+/+^ male mice beginning on P20. Data presented as mean ± SEM. Group numbers: Eu,*Dyrk1a*^+/Dox-Cre^ 15 F, 12 M; Eu,*Dyrk1a*^+/+^ 39 F, 32 M; Ts,*Dyrk1a*^+/+/+^ 13 F, 19 M; Ts,*Dyrk1a*^+/+/Dox-Cre^ 8 F, 14 M. ***p < 0.001, Eu,*Dyrk1a*^+/+^ > Ts,*Dyrk1a*^+/+/+^; ^#^p < 0.05, Eu,*Dyrk1a*^+/Dox-Cre^ < Eu,*Dyrk1a*^+/+^

### Ambulation

Female and male Ts,*Dyrk1a*^+/+/+^ mice achieved all measures of forward movement development significantly later than Eu,*Dyrk1a*^+*/*+^ mice (Fig. 2), and large effects were present in both sexes. The mean differences (in days) and Cohen’s effect sizes (**d**), respectively, were as follows: [females] Pivot: 1.4, **d**=.82; Crawl: 2.2, **d**=1.14; Transition: 2.2, **d**=1.20; Walk: 1.9, **d**=1.04; Run: 2.9, **d**=1.41; [males] Pivot: 2.1, **d**=1.48; Crawl: 2.2, **d**=1.38; Transition: 1.9, **d**=.97; Walk: 1.4, **d**=1.04; Run: 1.6, **d**=.72. The only significant improvement produced by functional reduction of *Dyrk1a* in trisomic mice was earlier achievement of the running milestone for male Ts,*Dyrk1a*^+/+/Dox-Cre^ mice as compared to Ts,*Dyrk1a*^+/+/+^ males (mean difference 1.3 days, **d**=.63). Otherwise, no improvements in forward movement were found in male or female Ts,*Dyrk1a*^+/+/Dox-Cre^ mice. Also, no significant differences between Eu,*Dyrk1a*^+*/*+^ and Eu,*Dyrk1a*^+/Dox-Cre^ mice were seen in either sex.

**Figure 2:**
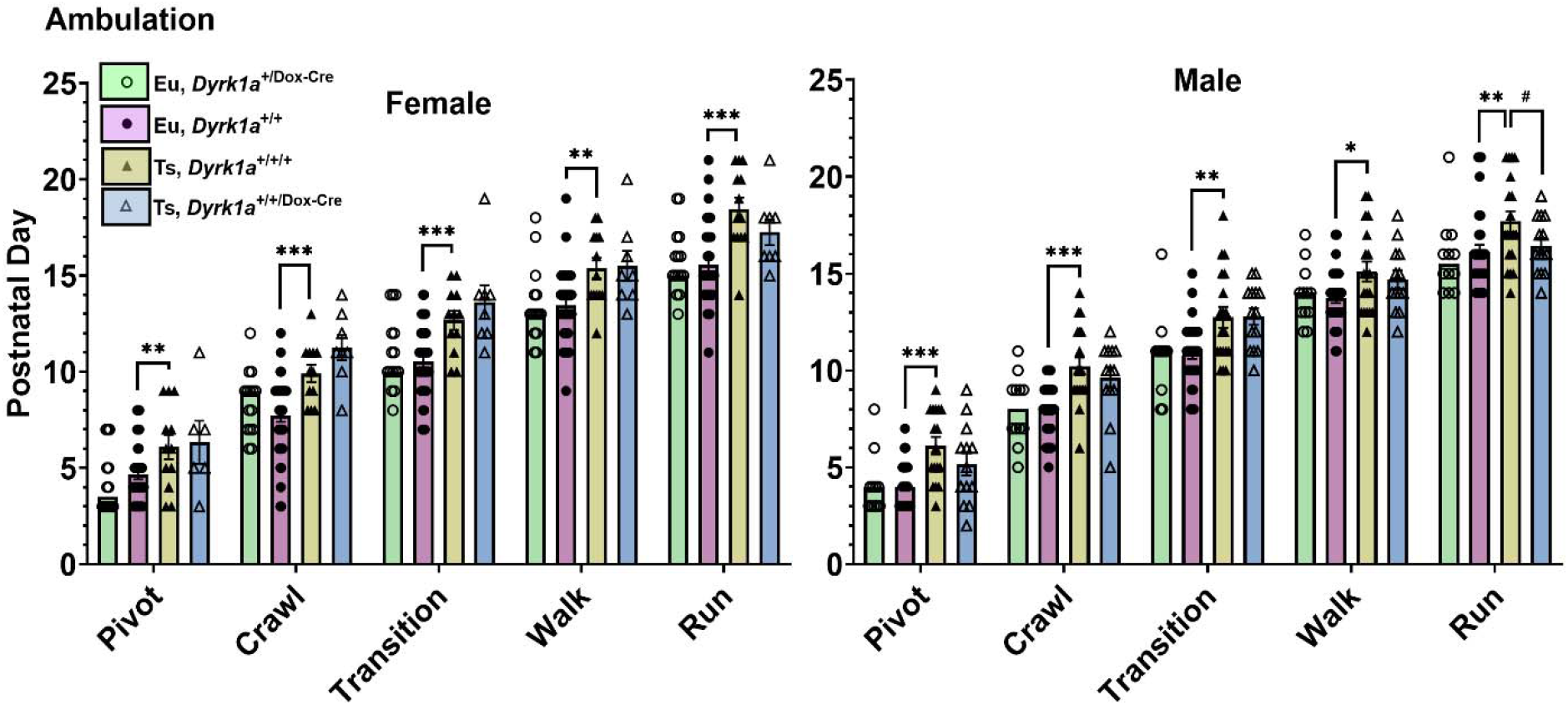
Forward Movement Development of Female and Male Pups. Female and male Ts,*Dyrk1a*^+/+/+^ mice were significantly delayed in achieving criteria for all five movement milestones as compared to Eu,*Dyrk1a*^+*/*+^ mice. The only significant improvement evident in Ts,*Dyrk1a*^+/+/Dox-Cre^ mice as compared to Ts,*Dyrk1a*^+/+/+^ littermates was for the running milestone for males. There were no significant differences between Eu,*Dyrk1a*^+/+^ and Eu,*Dyrk1a*^+/Dox-Cre^ mice. Data presented as mean ± SEM. Group numbers: Eu,*Dyrk1a*^+/Dox-Cre^ 15 F, 12 M; Eu,*Dyrk1a*^+/+^ 39 F, 32 M; Ts,*Dyrk1a*^+/+/+^ 13 F, 19 M; Ts,*Dyrk1a*^+/+/Dox-Cre^ 8 F, 14 M. *p < 0.05; **p < 0.01; ***p < 0.001, Ts,*Dyrk1a*^+/+/+^ > Eu,*Dyrk1a*^+*/*+^ #p < 0.05, Ts,*Dyrk1a*^+/+/Dox-Cre^ < Ts,*Dyrk1a*^+/+/+^

### Developmental Milestone Achievement

Euploid female and male pups achieved developmental milestones on approximately the same day, and trisomic pups showed delayed achievement of multiple milestones as compared to same sex euploid littermates (Fig. 3A). For females, Ts,*Dyrk1a*^+/+/+^ pups as compared to Eu,*Dyrk1a*^+/+^ littermates were significantly delayed in achieving all physical milestones except pinnae detachment and were significantly delayed in all sensorimotor milestones except the Preyer reflex. The mean differences (in days) and effect sizes (Cohen’s **d**) of the delayed milestones for females were as follows: lower incisor eruption: 0.9, **d**=0.67; upper incisor eruption; 0.9, **d**=0.81; eye opening: 1.5, **d**=1.31; cliff aversion: 5.1, **d**=1.86, negative geotaxis, 2.8, **d**=1.29; contact righting: 2.5, **d**=1.28; tactile stimulation: 2.6, **d**=1.37. For males, Ts,*Dyrk1a*^+/+/+^ pups were significantly delayed as compared to Eu,*Dyrk1a*^+/+^ littermates in two physical milestones (upper incisor eruption and eye opening) and in all five sensorimotor tasks. The mean differences (in days) and effect sizes (Cohen’s **d**) of the delayed milestones for males were as follows: upper incisor eruption; 0.5, **d**=0.50; eye opening: 1.5, **d**=1.15; cliff aversion: 5.3, **d**=1.84, negative geotaxis, 1.4, **d**=0.69; Preyer reflex: 0.9, **d**=0.72; contact righting: 1.3, **d**=0.55; tactile stimulation: 1.7, **d**=1.05. Although Ts,*Dyrk1a*^+/+/+^ animals of both sexes showed significant delays in most of the sensorimotor tests, for contact righting, tactile stimulation, and negative geotaxis the trisomic females as compared to trisomic males showed non-significant trends for longer delays (∼1 day longer) with larger effect sizes. Notably for cliff aversion, both female and male Ts,*Dyrk1a*^+/+/+^ mice had comparable and very long delays (>5 days) with very large effect sizes (**d**’s>1.8), indicating cliff aversion is a very sensitive measure of sensorimotor neurodevelopmental delays in this DS mouse model. Despite the sensitivity of these physical and sensorimotor measures to neurodevelopmental delays in the Ts65Dn mice, functional reduction of one copy of *Dyrk1a* in otherwise trisomic mice failed to produce significant improvement of neurodevelopmental delays in Ts,*Dyrk1a*^+/+/Dox-Cre^ mice as compared to Ts,*Dyrk1a*^+/+/+^ littermates in either sex (Fig. 3B). In addition, functional reduction of *Dyrk1a* in euploid mice yielded no significant differences between Eu,*Dyrk1a*^+*/*+^ and Eu,*Dyrk1a*^+/Dox-Cre^ mice in either sex (Fig. 3C).

**Figure 3:**
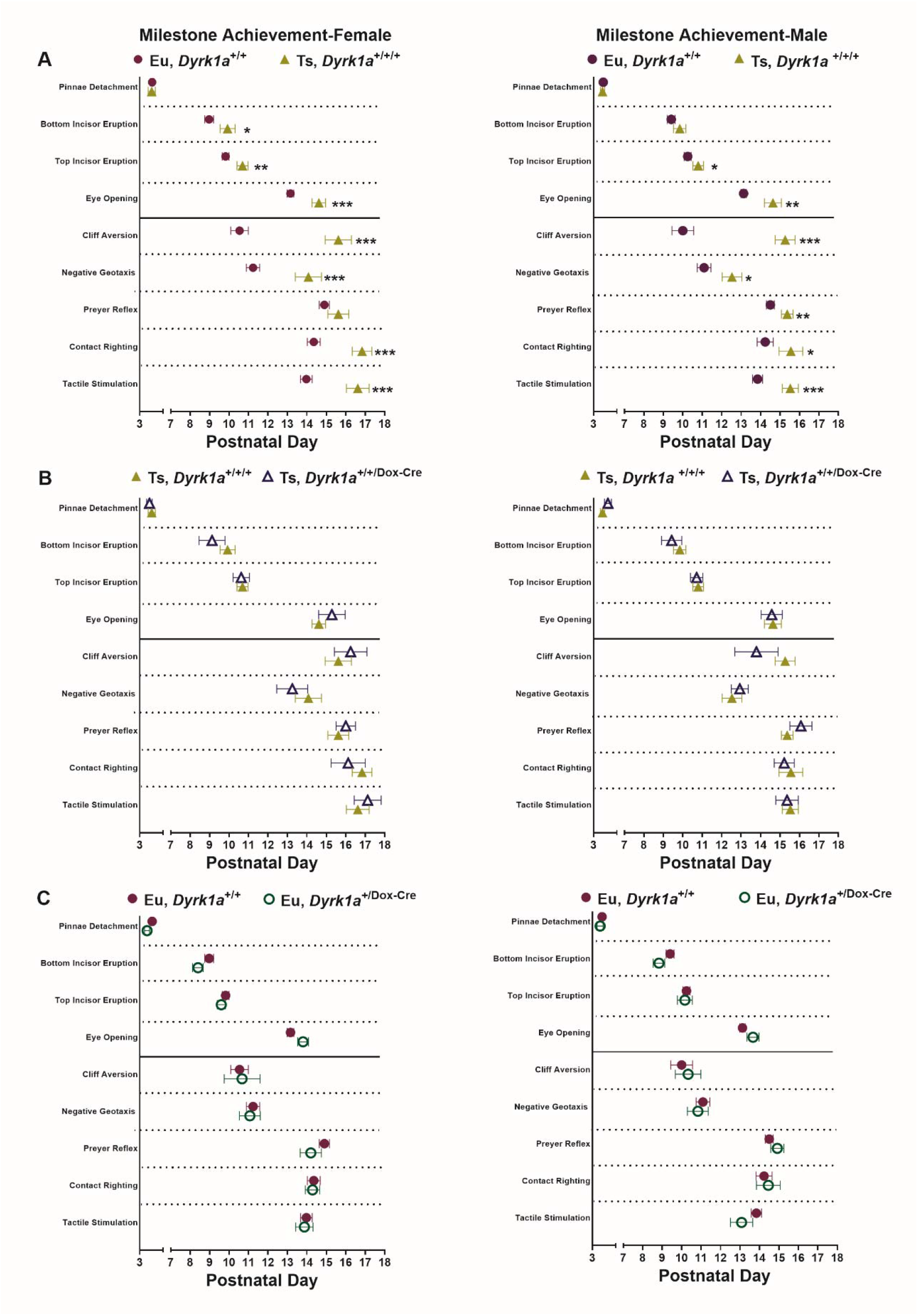
Female and Male Age of Achievement of Physical and Sensorimotor Milestones. (A) Female Ts,*Dyrk1a*^+/+/+^ as compared to Eu,*Dyrk1a*^+/+^ mice were significantly delayed in achieving all physical milestones except pinnae detachment. Trisomic females also were significantly delayed in all sensorimotor milestones except Preyer reflex. For male Ts,*Dyrk1a*^+/+/+^ as compared to Eu,*Dyrk1a*^+/+^ mice, upper incisor eruption and eye opening were significantly delayed among the physical milestones, whereas all sensorimotor milestones were significantly delayed. (B) There were no significant differences in milestone achievements in either sex for the Ts,*Dyrk1a*^+/+/Dox-Cre^ as compared to Ts,*Dyrk1a*^+/+/+^ littermates. (C) There were no significant differences for either sex between Eu,*Dyrk1a*^+/Dox-Cre^ and Eu,*Dyrk1a*^+/+^ littermates. Data presented as mean ± SEM. Group numbers: Eu,*Dyrk1a*^+/Dox-Cre^ 15 F, 12 M; Eu,*Dyrk1a*^+/+^ 39 F, 32 M; Ts,*Dyrk1a*^+/+/+^ 13 F, 19 M; Ts,*Dyrk1a*^+/+/Dox-Cre^ 8 F, 14 M *p < 0.05, **p < 0.01; ***p < 0.001 Ts,*Dyrk1a*^+/+/+^ > Eu,*Dyrk1a*^+/+^

### Ultrasonic Vocalization (USV)

#### Comparison of Ts,*Dyrk1a*^+/+/+^ and Eu,*Dyrk1a*^+*/*+^ mice

Trisomic as compared to euploid mice displayed fewer emitted calls in the first postnatal week but more calls (and a later decline) in the second postnatal week (Fig. 4A). These outcomes were consistent with the previous findings that first demonstrated a developmental shift in the peak rate of isolation-induced USVs in Ts65Dn mice from P6 to P9 [25]. In the first postnatal week, Eu,*Dyrk1a*^+*/*+^ pups emitted significantly more calls as compared to same-sex Ts,*Dyrk1a*^+/+/+^ littermates on P3 and P6 for females and on P3 for males. Additionally, in the first postnatal week euploid mice had longer call durations (Fig. 4B) and greater call power (Fig. 4C) than trisomic mice on P3 and P6. In contrast, in the second postnatal week Ts,*Dyrk1a*^+/+/+^ pups increased their rate of USV calls from P6 to P9 whereas Eu,*Dyrk1a*^+*/*+^ pups reduced their rate of USV calls from P6 to P12, resulting in significantly more calls by trisomic mice as compared to same-sex euploid littermates on P12 in females and on P9 and P12 in males (Fig. 4A). At all testing ages, the principal frequency (Fig. 4D) and peak frequency (data not shown) were lower in Eu,*Dyrk1a*^+*/*+^ as compared to Ts,*Dyrk1a*^+/+/+^ mice in both sexes.

**Figure 4:**
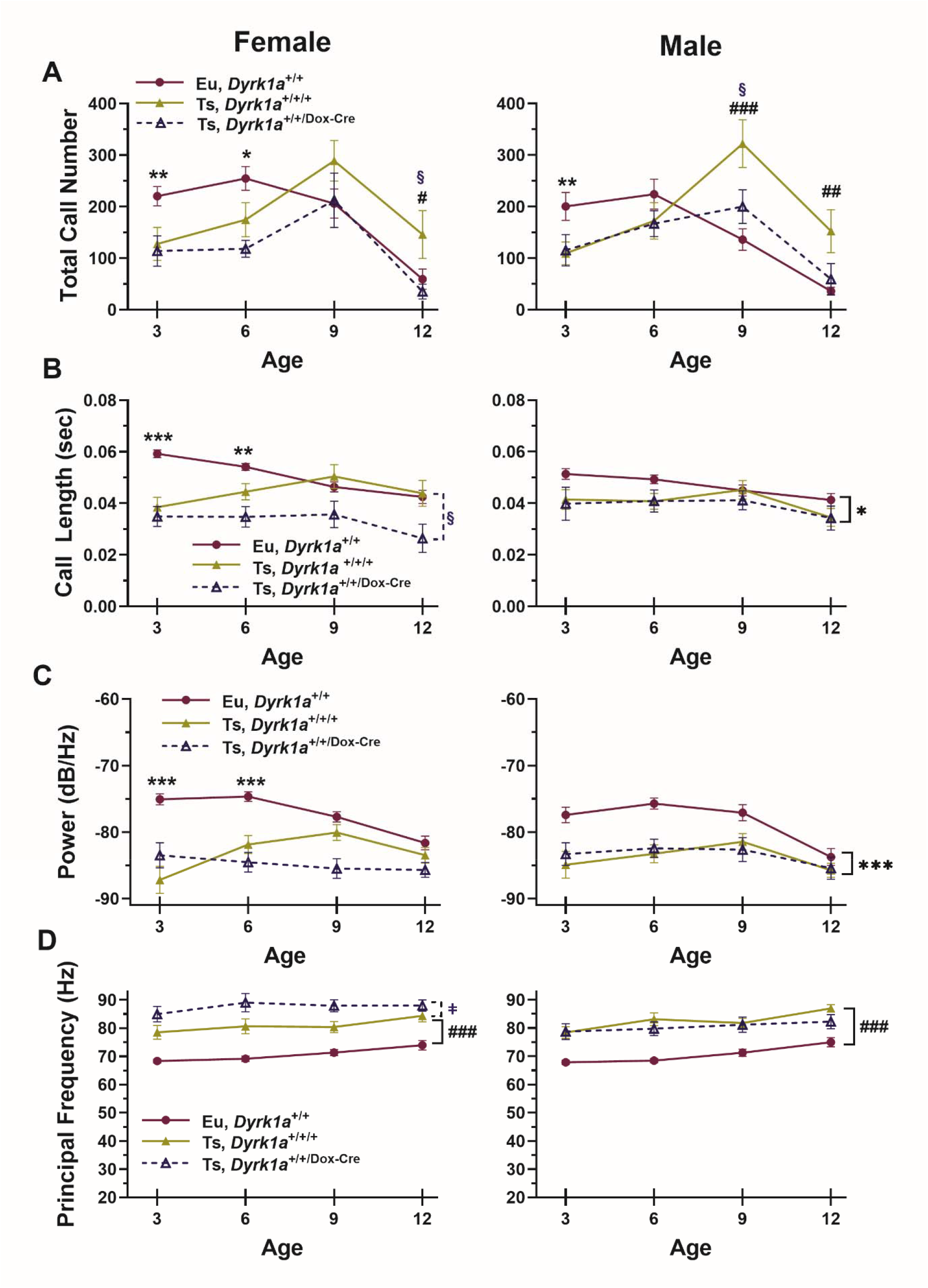
Isolation-induced Ultrasonic Vocalization Measures in Euploid (Eu,*Dyrk1a*^+/+^), Trisomic (Ts,*Dyrk1a*^+/+/+^), and Trisomic knockdown (Ts,*Dyrk1a*^+/+/Dox-Cre^) mice on Postnatal (P) Days 3, 6, 9, and 12. Data are presented as mean ± SEM. <u>(A)</u> *<u>Call Number</u>* Eu,*Dyrk1a*^+/+^ vs. Ts,*Dyrk1a*^+/+/+^: Genotype effects on total number of USV calls depended on age [Genotype × Age interaction, p<.01 for females, p<.001 for males]. Female and male Ts,*Dyrk1a*^+/+/+^ mice emitted fewer calls as compared to Eu,*Dyrk1a*^+/+^ mice in the first postnatal week (P3 and P6 for females; P3 for males [directional 1-tailed t-tests]), In contrast, the second postnatal week Ts,*Dyrk1a*^+/+/+^ pups emitted more calls than Eu,*Dyrk1a*^+/+^ pups (P12 in females; P9 and P12 in males [directional 1-tailed t-tests], confirming our a priori hypotheses. Ts,*Dyrk1a*^+/+/Dox-Cre^ vs. Ts,*Dyrk1a*^+/+/+^: There were no significant differences between Ts,*Dyrk1a*^+/+/Dox-Cre^ vs. Ts,*Dyrk1a*^+/+/+^ in the first postnatal week. In the second postnatal week, Ts,*Dyrk1a*^+/+/Dox-Cre^ mice emitted significantly fewer USV calls as compared to Ts,*Dyrk1a*^+/+/+^ mice (P12 in females; P9 in males [directional 1-tailed t-tests]), approaching the values of the euploid controls in the second week. <u>(B)</u> *<u>Call Length</u>* Eu,*Dyrk1a*^+/+^ vs. Ts,*Dyrk1a*^+/+/+^: For females, the average call length was significantly reduced in Ts,*Dyrk1a*^+/+/+^ mice as compared to Eu,*Dyrk1a*^+/+^ mice on P3 and P6 [Genotype × Age interaction, p<.01; post hoc comparisons at each age used 2-tailed t-tests]. For males, average call length was reduced across days in Ts,*Dyrk1a*^+/+/+^ as compared to Eu,*Dyrk1a*^+/+^ mice [main effect of Genotype, p<.05]. Ts,*Dyrk1a*^+/+/Dox-Cre^ vs. Ts,*Dyrk1a*^+/+/+^: For females, the average call length across age was significantly shorter in Ts,*Dyrk1a*^+/+/Dox-Cre^ mice as compared to Ts,*Dyrk1a*^+/+/+^ [main effect of Genotype, p<.05). For males, there were no significant differences between Ts,*Dyrk1a*^+/+/Dox-Cre^ and Ts,*Dyrk1a*^+/+/+^ mice. <u>(C)</u> *<u>Call Power</u>* Eu,*Dyrk1a*^+/+^ vs. Ts,*Dyrk1a*^+/+/+^: Average call power (in dB/Hz) for female Ts,*Dyrk1a*^+/+/+^ mice was reduced compared to Eu,*Dyrk1a*^+/+^ females on P3 and P6 [Genotype × Age interaction (p<.001); post hoc 2-tailed t-tests]. For males, call power was reduced across days in Ts,*Dyrk1a*^+/+/+^ as compared to Eu,*Dyrk1a*^+/+^ mice [Genotype main effect (p<.001)]. Ts,*Dyrk1a*^+/+/Dox-Cre^ vs. Ts,*Dyrk1a*^+/+/+^: For average call power in the Ts,*Dyrk1a*^+/+/+^ and Ts,*Dyrk1a*^+/+/Dox-Cre^ analysis, there were no significant main or interactive effects of genotype in either sex. <u>(D)</u> *<u>Principal Frequency</u>* Eu,*Dyrk1a*^+/+^ vs. Ts,*Dyrk1a*^+/+/+^: The average principal frequency (in Hz) was higher in Ts,*Dyrk1a*^+/+/+^ as compared to Eu,*Dyrk1a*^+/+^ mice in females and in males across all four ages [main effect of Genotype in each, p<.001)]. Similar outcomes were present in *Peak Frequency* (data not shown). Ts,*Dyrk1a*^+/+/Dox-Cre^ vs. Ts,*Dyrk1a*^+/+/+^: In females, the average principal frequency was higher in Ts,*Dyrk1a*^+/+/Dox-Cre^ mice as compared to Ts,*Dyrk1a*^+/+/+^ mice [main effect of Genotype, p<.01)]. In males, there were no significant differences between the Ts,*Dyrk1a*^+/+/Dox-Cre^ as compared to Ts,*Dyrk1a*^+/+/+^ mice. Similar outcomes were seen for females and males for *Peak Frequency* (data not shown). Eu,*Dyrk1a*^+/+^ (n=36-39 F, 28-30 M) vs. Ts,*Dyrk1a*^+/+/+^ (n=13F, 18-19M) *p < 0.05; **p < 0.01; ***p < 0.001, Eu,*Dyrk1a*^+*/*+^ > Ts,*Dyrk1a*^+/+/+^ #p < 0.05; ^##^p < 0.01; ^###^p < 0.001, Ts,*Dyrk1a*^+/+/+^ > Eu,*Dyrk1a*^+*/*+^ {solid black right bracket} Main effect of Genotype (interaction with Age was not significant) Ts,*Dyrk1a*^+/+/+^ (n=13F, 18-19M) vs. Ts,*Dyrk1a*^+/+/Dox-Cre^ (n=8F, 9M) **^§^**p < 0.05; Ts,*Dyrk1a*^+/+/+^ > Ts,*Dyrk1a*^+/+/Dox-Cre^ □p < 0.05; Ts,*Dyrk1a*^+/+/+^ < Ts,*Dyrk1a*^+/+/Dox-Cre^ 4

#### Comparison of Ts,*Dyrk1a*^+/+/+^ and Ts,*Dyrk1a*^+/+/Dox-Cre^ mice

The a priori hypothesis that normalizing *Dyrk1a* copy number in otherwise trisomic mice (Ts,*Dyrk1a*^+/+/Dox-Cre^) would improve the developmental trajectory of USVs as compared to trisomic mice (Ts,*Dyrk1a*^+/+/+^) received partial support. During the second postnatal week (Fig. 4A), female Ts,*Dyrk1a*^+/+/Dox-Cre^ emitted significantly fewer USVs than female Ts,*Dyrk1a*^+/+/+^ mice on P12, and male Ts,*Dyrk1a*^+/+/Dox-Cre^ emitted significantly fewer calls on P9 with a strong trend for fewer calls on P12 (p=.051). In contrast, USV call number in the first postnatal week did not differ significantly between the Ts,*Dyrk1a*^+/+/Dox-Cre^ and the Ts,*Dyrk1a*^+/+/+^ groups in either sex. Additionally, for males there were no significant effects of genotype on call length, call power, or principal frequency (Fig. 4 B,C,D). For females, across the four ages there was a main effect of genotype reflecting the significantly shorter call lengths (p<.05, Fig. 4B) and significantly higher average principal frequency (p<.01, Fig. 4D) and average peak frequency (p<.05, data not shown) of the Ts,*Dyrk1a*^+/+/Dox-Cre^ mice as compared to the Ts,*Dyrk1a*^+/+/+^ mice. Note that for these effects on call length and call frequency, the direction of effects in the Ts,*Dyrk1a*^+/+/Dox-Cre^ mice made them more divergent from euploids than the Ts,*Dyrk1a*^+/+/+^ mice. In summary, the diminished number of calls over the second postnatal week was consistent with improved developmental USV phenotypes following functional reduction of *Dyrk1a* in otherwise trisomic mice.

#### Comparison of Eu,*Dyrk1a*^+*/*Dox-Cre^ and Eu,*Dyrk1a*^+*/*+^

Functional reduction of one copy of *Dyrk1a* in otherwise euploid mice (Eu,*Dyrk1a*^+*/*Dox-Cre^) did not produce any significant effects as compared to euploid littermates (Eu,*Dyrk1a*^+*/*+^) in either sex on any USV measure at any test age (Supplemental Fig. 6). Eu,*Dyrk1a*^+*/*+^ and Eu,*Dyrk1a*^+*/*Dox-Cre^ mice both displayed the expected and similar significant reductions in USV call number over days (p’s<.001), along with similar reductions in call length and call power over days (p’s<.001).

#### Body Temperature Effects During USV Testing

All groups showed large reductions in body temperature on the P3 post-USV measurement compared to the pre-USV measurement, and the temperature drops became progressively less severe from P6 to P12. For the first directional a priori hypothesis (Fig. 5), Ts,*Dyrk1a*^+/+/+^ mice of both sexes had significantly more severe body temperature reductions than Eu,*Dyrk1a*^+/+^ littermates on P6, P9, and P12. These effects confirm a significant developmental delay in the Ts,*Dyrk1a*^+/+/+^ mice in the emergence of thermoregulatory responses to isolation at room temperature.

**Figure 5.**
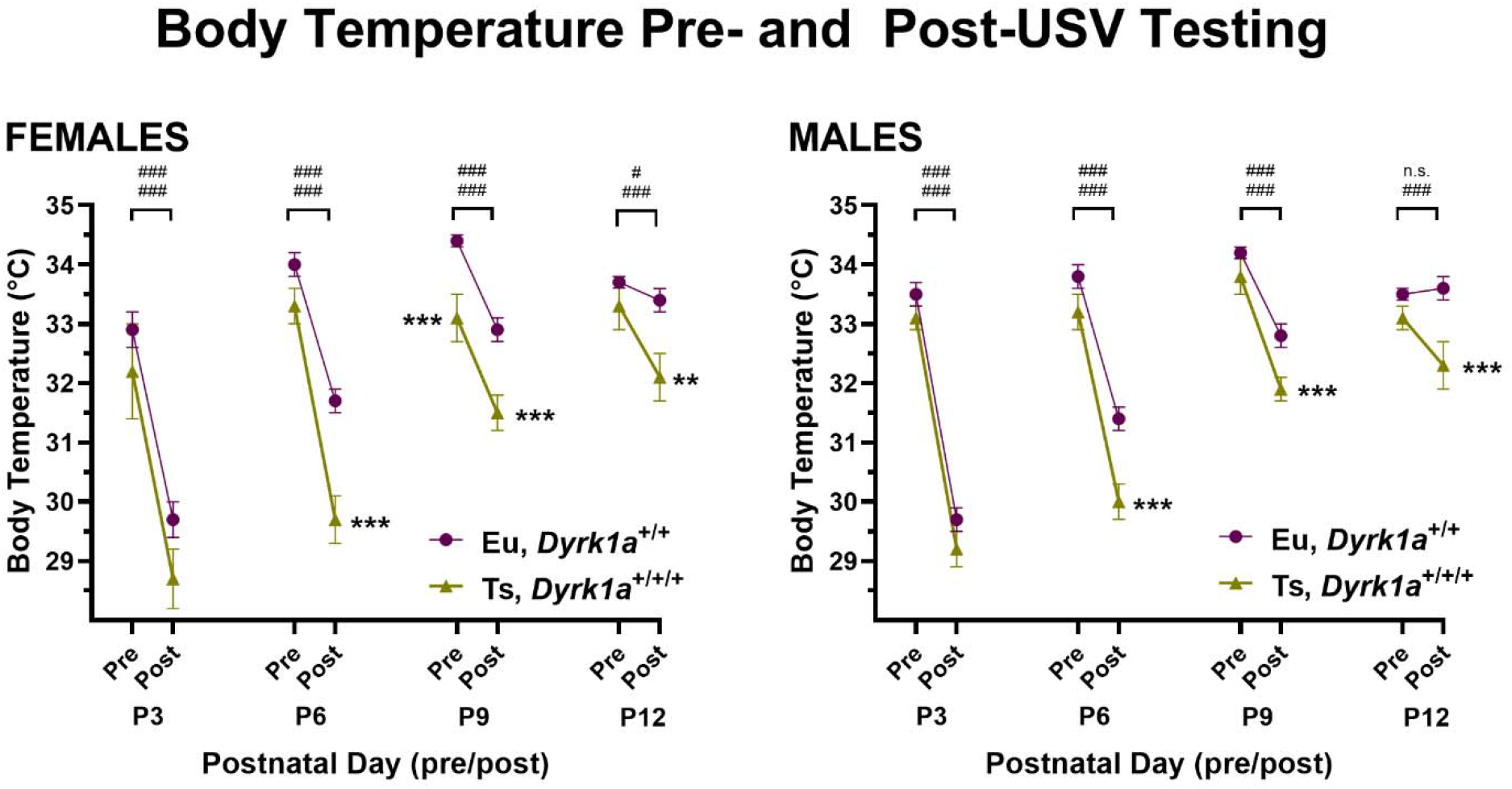
Body Temperatures Before and After Each USV Test Session on P3, P6, P9, and P12 for Female and Male Mice of Eu,*Dyrk1a*^+/+^ and Ts,*Dyrk1a*^+/+/+^ Genotypes. All groups showed significant body temperature loss on P3 that became progressively less severe with increasing age [main effects of day and pre/post and day × pre/post interactions were significant, p’s<.001]. For the directional a priori hypothesis of greater isolation-induced temperature reductions in Ts,*Dyrk1a*^+/+/+^ mice than Eu,*Dyrk1a*^+/+^mice, both female and male Ts,*Dyrk1a*^+/+/+^ had significantly more severe body temperature reductions than Eu,*Dyrk1a*^+/+^ littermates on P6, P9, and P12 [RM-ANOVA genotype × pre/post interaction (p<.001), main effect of genotype (p<.001), and post-hoc Sidek comparisons as shown]. ^#^p<.05; ^###^p<.001, Sidek post hoc tests: the Post-USV test was significantly reduced relative to the Pre-USV test for the session; n.s.=not significant (note: the top and bottom symbols refer to the top and bottom groups listed in the panel legend, respectively) *p<.05; **p<.01; ***p<.001, Sidek post hoc tests, Ts,*Dyrk1a*^+/+/+^ group was significantly lower than comparison group

For the second directional a priori hypothesis [Ts,*Dyrk1a*^+/+/Dox-Cre^ < Ts,*Dyrk1a*^+/+/+^, Supplemental Fig.7A], there were no significant main or interactive effects of genotype for either sex, indicating that normalization of *Dyrk1a* copy number did not mitigate the developmental deficits in response to thermoregulatory challenge in trisomic mice. For the third a priori hypothesis [Eu,*Dyrk1a*^+/Dox-Cre^ ≠ Eu,*Dyrk1a*^+/+^, Supplemental Fig. 7B], females yielded a significant 3-way interaction that was due mainly to the Eu,*Dyrk1a*^+/Dox-Cre^ group having a significantly higher body temperature on the P6 post-test measurement and a trend for a higher body temperature on the P3 the pre-test measurement. For males, a significant main effect of genotype was due mainly to the higher body temperatures of the Eu,*Dyrk1a*^+/Dox-Cre^ group as compared to Eu,*Dyrk1a*^+/+^ group for the pre-test measurement on P3 and P12. These data suggest that *Dyrk1a* haploinsufficiency in otherwise euploid mice may modestly accelerate thermoregulatory development in early postnatal pups.

**Figure 6:**
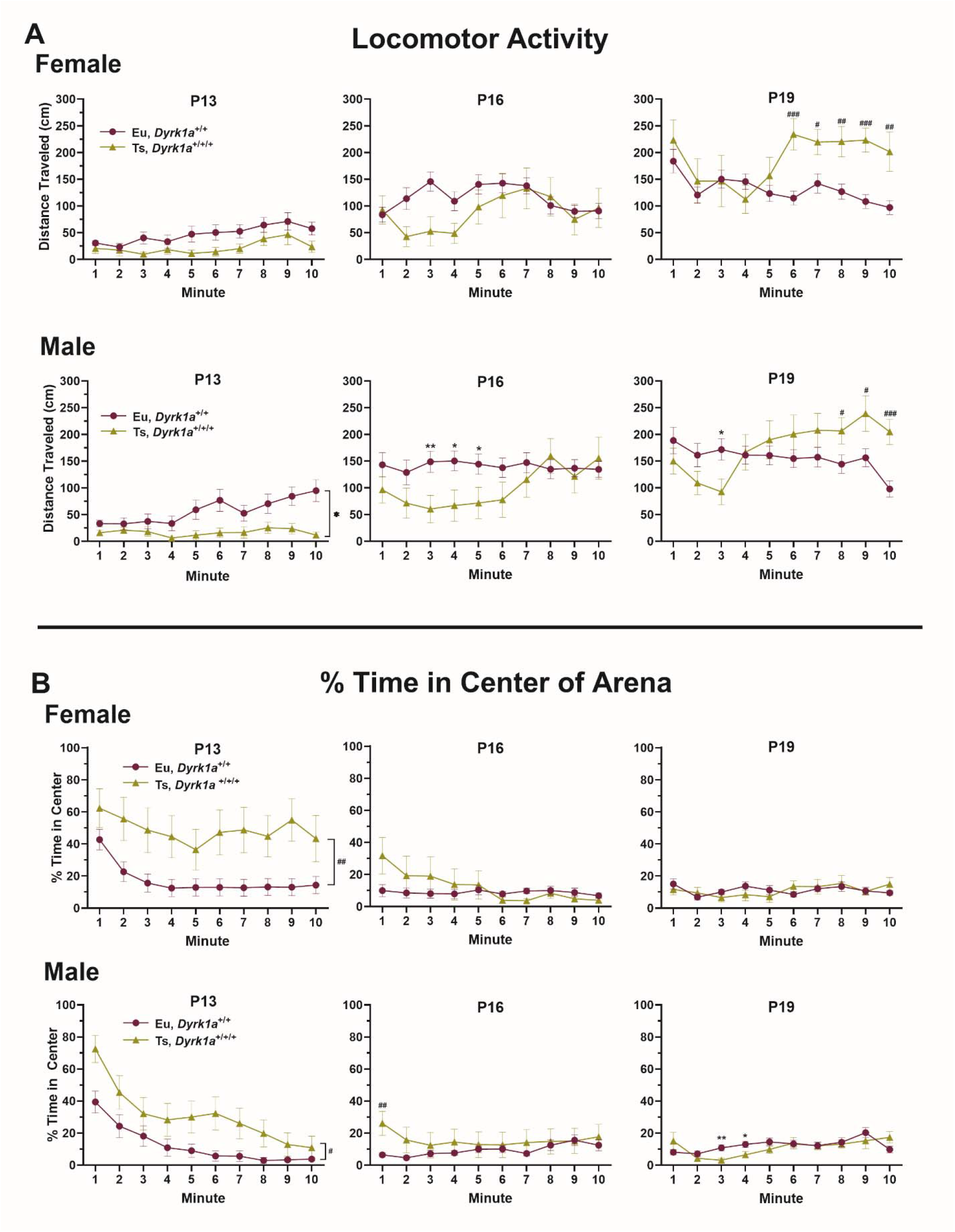
Distance Traveled (A) and Percent Time Spent in the Center of the Arena (B) during Locomotor Activity Test in Female and Male Euploid (Eu,*Dyrk1a*^+/+^) and Trisomic (Ts,*Dyrk1a*^+/+/+^) mice on Postnatal (P) Days 13, 16, and 19. Data presented as mean ± SEM. Mice of both genotypes showed increases in locomotor activity over development, and the elevated activity predicted for the trisomic mice emerged at P19. On P13, overall activity levels were relatively low yet Ts,*Dyrk1a*^+/+/+^ males were significantly less active than Eu,*Dyrk1a*^+/+^ males (main effect of genotype, p<.05); the trend for lower activity in Ts,*Dyrk1a*^+/+/+^ females at P13 was not significant. On P16, the Ts,*Dyrk1a*^+/+/+^ males were less active than Eu,*Dyrk1a*^+/+^ males in the first half of the session (Genotype × Time interaction, p<.05) and were significantly lower for minutes 3-5; a similar trend in the Ts,*Dyrk1a*^+/+/+^ females on P16 did not reach significance. On P19, Ts,*Dyrk1a*^+/+/+^ mice of both sexes showed significant elevations in locomotor activity in the second half of the session (Genotype × Time interaction, females p<.01; males p<.001), with significantly longer distances traveled in each of the last 5 minutes in females and each of the last 3 minutes in males. Trisomic males were also significantly less active than euploid males in minute 3 of the session. Mice of both genotypes also showed reductions over development. On P13, the Ts,*Dyrk1a*^+/+/+^ mice of both sexes spent significantly more time in the center than euploid littermates. On P16, Ts,*Dyrk1a*^+/+/+^ male mice spent significantly more time in the center in the first minute (and females showed a similar non-significant trend), but for the remainder of the session time in center was low and no differences were seen between genotypes. On P19, mice of both groups in both sexes spent <20% in the center on average for all groups, even including the small but significant difference in time in center for male Eu,*Dyrk1a*^+/+^ and Ts,*Dyrk1a*^+/+/+^ mice during minutes 3-4 Group numbers: Eu,*Dyrk1a*^+/+^ 36-37 F, 31 M; Ts,*Dyrk1a*^+/+/+^ 11-12F, 17-19 M *p < 0.05; **p < 0.01, Eu,*Dyrk1a*^+/+^ > Ts,*Dyrk1a*^+/+/+^ #p < 0.05; ^##^p < 0.01; ^###^p < 0.001, Ts,*Dyrk1a*^+/+/+^ > Eu,*Dyrk1a*^+*/*+^ Right brackets: Main effect of Genotype (interaction with Age was not significant)

**Figure 7:**
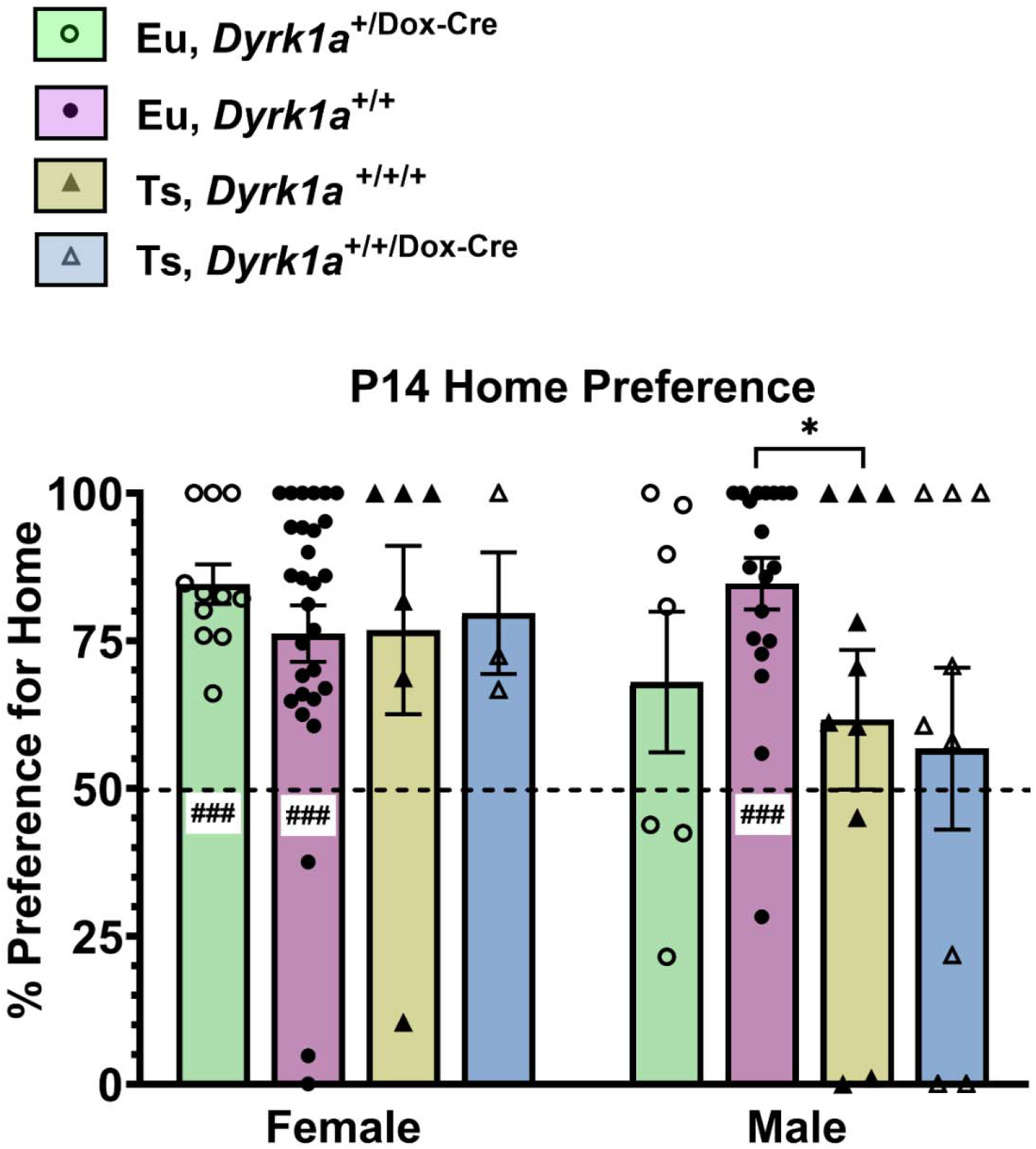
Home Bedding Preference Ratio on P14. Home preference ratio was calculated using the equation: (time in home zone)/(time in home zone + time in clean zone). Tests of preference scores against chance (50%) for each group using one-sample ttests indicated female and male Eu,*Dyrk1a*^+/+^ pups and female Eu,*Dyrk1a*^+/Dox-Cre^ pups were significantly above chance. A priori comparisons between Ts,*Dyrk1a*^+/+/+^ and Eu,*Dyrk1a*^+/+^ pups showed confirmed significantly lower home preference in males but not in females. No other a priori comparisons reached significance. Data presented as mean ± SEM. Group numbers: Eu,*Dyrk1a*+/Dox-Cre n=11 F, 7 M; Eu,*Dyrk1a*^+/+^ n=28 F, 19 M; Ts,*Dyrk1a*^+/+/+^ n=6 F, 10 M; Ts,*Dyrk1a*^+/+/Dox-Cre^ n=3 F, 9 M. *p < 0.05, Eu,*Dyrk1a*^+*/*+^ > Ts,*Dyrk1a*^+/+/+^, a priori directional 1-tailed t-test ^###^p < 0.001, group significantly greater than chance, one-sample t-test

### Locomotor Activity

#### Comparison of P10 Groups of mice

Locomotor activity on P10 was limited (overall median of total distance traveled: 76 cm for females; 46 cm for males) and from the fifth minute onward, activity never exceeded 10 cm per minute for any group. Consequently, only total activity and overall percent time in the center were analyzed for the P10 session (Supplemental Table 3). Eu,*Dyrk1a*^+/+^ females traveled significantly farther (p<.001) and spent significantly less time in the center (p<.001) as compared to Ts,*Dyrk1a*^+/+/+^ females. For males, the Eu,*Dyrk1a*^+/+^ pups also spent significantly less time in the center than male Ts,*Dyrk1a*^+/+/+^ littermates (p=.022), but the trend for male euploid mice for greater distance traveled than trisomic mice did not reach significance (p=.083). Functional reduction of one copy of *Dyrk1a* did not alter outcomes on these locomotor activity tests on P10 either in trisomic mice or in euploid mice, with no significant differences either between Ts,*Dyrk1a*^+/+*/*+^ and Ts,*Dyrk1a*^+/+*/*Dox-Cre^ groups or between Eu,*Dyrk1a*^+/+^ and Eu,*Dyrk1a*^+*/*Dox-Cre^ groups for either sex.

#### P13 to P19: Comparison of Eu,*Dyrk1a*^+/+^ and Ts,*Dyrk1a*^+/+/+^

Mice of both genotypes showed an increase in locomotor activity across age in both sexes, and hyperactivity predicted for the trisomic mice as compared to euploid littermates emerged at P19 (Fig. 6A). On P13, despite the relatively low overall activity levels in all groups, the distance traveled by Ts,*Dyrk1a*^+/+/+^ males was significantly less than Eu,*Dyrk1a*^+/+^ males (main effect of genotype, p<.05); female Ts,*Dyrk1a*^+/+/+^ mice exhibited a similar but non-significant trend for lower activity. On P16, Ts,*Dyrk1a*^+/+/+^ males were significantly less active than Eu,*Dyrk1a*^+/+^ males in the first half of the session (Genotype × Time interaction, p<.05) with significant reductions in distance traveled during minutes 3-5. A similar trend in the Ts,*Dyrk1a*^+/+/+^ females on P16 did not reach significance. In contrast, on P19 the Ts,*Dyrk1a*^+/+/+^ mice of both sexes showed significant elevations in locomotor activity in the second half of the session (Genotype × Time interaction, females p<.01; males p<.001), with significantly longer distances traveled in each of the last 5 minutes in females and in each of the last 3 minutes in males. The elevated activity in the last 3 minutes in males contrasted with their reduced activity earlier in the session (on minute 3).

Concurrent with the developmental increase in overall activity from P13 to P19, the time spent in the center declined sharply in both genotypes after P13 (Fig. 6B). On P13, female and male Ts,*Dyrk1a*^+/+/+^ mice spent significantly more time in the center than same-sex Eu,*Dyrk1a*^+/+^ littermates. On P16, differences between Ts,*Dyrk1a*^+/+/+^ and Eu,*Dyrk1a*^+/+^ were present only during the first minute, with Ts,*Dyrk1a*^+/+/+^ male mice spending significantly more time in the center than Eu,*Dyrk1a*^+/+^ males, with a similar but non-significant trend in females. After the first minute of the P16 session, time in center was low and no differences were seen between genotypes. On P19, mice of all groups in both sexes largely avoided the center zone, spending <20% of the time on average of each minute, though male Eu,*Dyrk1a*^+/+^ mice had small but significant increases compared to male Ts,*Dyrk1a*^+/+/+^ mice during minutes 3 and 4.

#### P13 to P19: Comparison of Ts,*Dyrk1a*^+/+/+^ and Ts,*Dyrk1a*^+/+/Dox-Cre^ mice

Locomotor activity increased across age, but no significant differences were exhibited between Ts,*Dyrk1a*^+/+/+^ and Ts,*Dyrk1a*^+/+/Dox-Cre^ mice for either sex at any age (Supplemental Fig. 8). For percent time in the center (Supplemental Fig. 9), there were no significant main or interactive effects of genotype for either sex on P13 (when time in center was relatively high) and the trend on P13 for greater time in center in the second half of the session for Ts,*Dyrk1a*^+/+/+^ female mice did not reach significance (Genotype × Minute interaction, p=.063). On P16, time in center was relatively low after the first minute and there were no significant main or interactive effects of genotype in females. In males, the greater reductions in the last four minutes in Ts,*Dyrk1a*^+/+/Dox-Cre^ mice as compared to Ts,*Dyrk1a*^+/+/+^ mice yielded a Genotype × Minute interaction (p=.033), but individual comparisons on minutes 7-10 did not reach significance. On P19 (when time in center was relatively low), there were no significant main or interactive effects of genotype for either sex. Overall, for both P16 and P19, the mice of all groups largely avoided the center zone after the first minute, with average time in center all <20% on the remaining 9 minutes.

#### P13 to P19: Comparison of Eu,*Dyrk1a*^+/+^ and Eu,*Dyrk1a*^+/Dox-Cre^ mice

For locomotor activity of Eu,*Dyrk1a*^+/+^ and Eu,*Dyrk1a*^+/Dox-Cre^ groups (Supplemental Fig. 10), there were no significant main or interactive effects of genotype on P13 (either sex), P16 (females), or P19 (either sex). The only group difference was in males on P16, when Eu,*Dyrk1a*^+/+^ males traveled significantly farther during the session than Eu,*Dyrk1a*^+/Dox-Cre^ males (main effect of genotype, p<.05). For percent time in center (Supplemental Fig. 11), there were no significant differences between Eu,*Dyrk1a*^+/+^ and Eu,*Dyrk1a*^+/Dox-Cre^ groups of either sex at any age.

### Homing Test

In the 1-minute homing test on P11 and P14, a sample of home bedding and a sample of clean bedding were placed at one end of the test cage and the time to reach each cup and the relative amount of time spent in the home bedding (as a preference ratio) was assessed. At P11, however, 72 of the 125 mice (56%) failed to traverse the test cage far enough to reach either cup, ranging across groups from 38% to 100% (see Supplemental Table 4). Moreover, 42 of the 125 mice (34%) failed even to cross the midline of the test cage. The limited mobility affecting all groups on P11 prevented valid analysis of homing performance at this age. Notably, however, the percentage of Ts,*Dyrk1a*^+/+/+^ mice (sexes combined) that failed to attain traversal benchmarks was significantly greater than Eu,*Dyrk1a*^+/+^ mice both for reaching the midline (52% vs. 19%, p<.01) and for reaching either cup (87% vs. 47% p<.001, Fishers Exact Test).

On P14, the majority of mice (93/119, 78%) traversed the test cage and entered at least one of the cups (Supplemental Table 5), with no significant differences among groups in the proportion of mice failing to reach the cups. Mice that reached the cups were included in the analysis of measures of preference for home bedding. As shown in Fig. 7, the home preference scores in the male and female Eu,*Dyrk1a*^+/+^ mice and female Eu,*Dyrk1a*^+/Dox-Cre^ mice were significantly higher than chance. Only male Ts,*Dyrk1a*^+/+/+^ pups showed deficits in home preference, showing significantly lower home preference than Eu,*Dyrk1a*^+/+^ males (p<.05). For the latency to reach the cups, there were no significant main or interactive effects of genotype in either sex in the three a priori Genotype × Cup Type repeated measures ANOVAs. Across all groups, females exhibited significantly shorter latencies to the home as compared to the clean cup [19.4 and 33.1 sec, respectively, p<.001], but the difference in males across all groups did not reach significance [25.8 and 34.2 sec, respectively, p=.115].

### DYRK1A Protein and mRNA Expression

DYRK1A protein and mRNA expression were quantified in the hippocampus, cerebral cortex and cerebellum tissues in Eu,*Dyrk1a*^+/+^, Eu,*Dyrk1a*^+/Dox-Cre^ , Ts,*Dyrk1a*^+/+/+^, and Ts,*Dyrk1a*^+/+/Dox-Cre^ animals at P6 to evaluate the short-term effects of gene copy number reduction, and at P21 to determine how *Dyrk1a* gene excision at birth affected the brain at the time of weaning (after testing). As shown in Fig. 8A, DYRK1A was significantly elevated in female Ts,*Dyrk1a*^+/+/+^ as compared to female Eu,*Dyrk1a*^+/+^ pups in the cerebral cortex, with trends for increased expression in the hippocampus and cerebellum. In male mice at P6, DYRK1A was significantly elevated in the hippocampus, cerebral cortex, and cerebellum of Ts,*Dyrk1a*^+/+/+^ pups as compared euploid littermate controls. The postnatal functional reduction of *Dyrk1a* copy number on P0 significantly reduced protein expression levels of P6 male Ts,*Dyrk1a^+/+^*^Dox-Cre^ pups as compared to Ts,*Dyrk1a*^+/+/+^ mice in the cerebellum, but not other regions in males or in any region in females. In mRNA from P6 mice (Fig. 8B), levels were significantly elevated in the hippocampus, cerebral cortex, and cerebellum of female and male Ts,*Dyrk1a*^+/+/+^ as compared to Eu,*Dyrk1a*^+/+^ animals. The postnatal functional reduction of *Dyrk1a* did not significantly alter mRNA expression on P6 either for Ts,*Dyrk1a^+/+^*^Dox-Cre^ compared to Ts,*Dyrk1a*^+/+/+^ groups or for Eu,*Dyrk1a*^+/Dox-Cre^ as compared to Eu,*Dyrk1a*^+/+^ groups.

**Figure 8:**
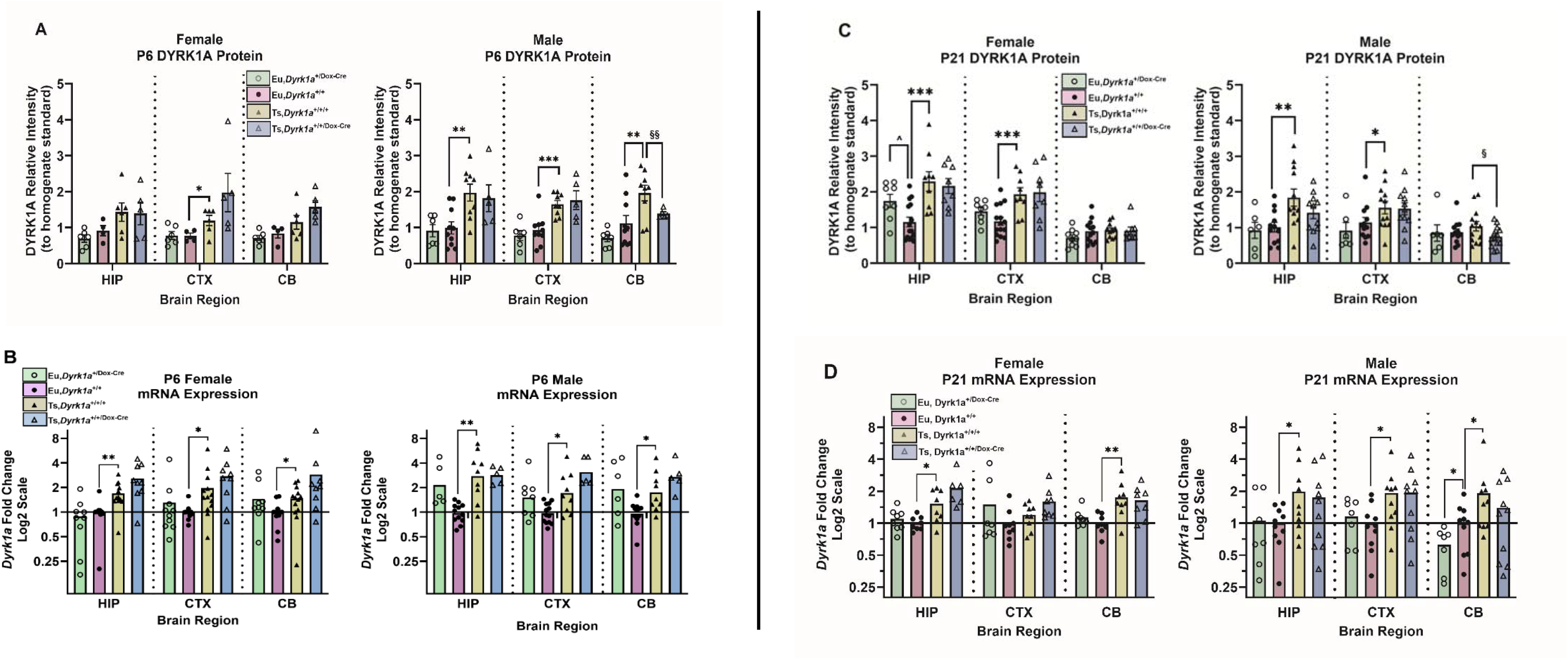
Female and Male DYRK1A Protein Expression on Postnatal (P) Day 6 (A) and P21 (C) and *Dyrk1a* mRNA Expression on P6 (B) and P21 (D). Protein data are presented as mean ± SEM relative to the within-blot protein homogenate mean; the mRNA data are presented as mean ± SEM relative ratios to within-plate euploid mean. **A:** As compared to Eu,*Dyrk1a*^+/+^ mice of the same sex, DYRK1A protein was significantly overexpressed in Ts,*Dyrk1a*^+/+/+^ males in the hippocampus (HIP), cerebral cortex (CTX), and cerebellum (CB), and in Ts,*Dyrk1a*^+/+/+^ females in the cerebral cortex. Functional reduction of one copy of *Dyrk1a* significantly reduced DYRK1A overexpression in the Ts,*Dyrk1a*^+/+/Dox-Cre^ male cerebellum as compared to Ts,*Dyrk1a*^+/+/+^ males. [Group numbers: Eu,*Dyrk1a*^+/Dox-Cre^ n=6 F, 6 M; Eu,*Dyrk1a*^+/+^ n=4 F, 10 M; Ts,*Dyrk1a*^+/+/+^ n=6 F, 9 M; Ts,*Dyrk1a*^+/+/Dox-Cre^ n=5 F; 5 M] B. *Dyrk1a* mRNA transcripts were overexpressed in the hippocampus (HIP), cerebral cortex (CTX), and cerebellum (CB) of Ts,*Dyrk1a*^+/+/+^ animals of both sexes. [Group numbers: Eu,*Dyrk1a*^+/Dox-Cre^ n=9-10 F, 5-9 M; Eu,*Dyrk1a*^+/+^ n=12 F, 12-13M; Ts,*Dyrk1a*^+/+/+^ n=12-13 F, 9M; Ts,*Dyrk1a*^+/+/Dox-Cre^ n=11 F, 5 M] C. As compared to Eu,*Dyrk1a*^+/+^ mice of the same sex, DYRK1A protein was significantly overexpressed in P21 female and male Ts,*Dyrk1a*^+/+/+^ hippocampus and cerebral cortex. In addition, DYRK1A protein was significantly reduced in male Ts,*Dyrk1a*^+/+/Dox-Cre^ cerebellum as compared to Ts,*Dyrk1a*^+/+/+^ males.[Group numbers: Eu,*Dyrk1a*^+/Dox-Cre^ n=8-9 F, 6 M; Eu,*Dyrk1a*^+/+^ n=13-15 F, 12-13 M; Ts,*Dyrk1a*^+/+/+^ n=9-11F, 12-13M; Ts,*Dyrk1a*^+/+/Dox-Cre^ n=8F, 11-13M] **D:** *Dyrk1a* mRNA transcripts were significantly overexpressed in P21 female hippocampus and cerebellum and in male Ts65Dn hippocampus, cerebral cortex, and cerebellum. Group numbers: Eu,*Dyrk1a*^+/Dox-Cre^ n=8 F, 7 M; Eu,*Dyrk1a*^+/+^ n=10 F, 10 M; Ts,*Dyrk1a*^+/+/+^ n=9-10 F, 10 M; Ts,*Dyrk1a*^+/+/Dox-Cre^ n=7-8 F, 10 M. *p < 0.05; **p < 0.01; ***p < 0.001: Ts,*Dyrk1a*^+/+/+^ > Eu,*Dyrk1a*^+*/*+^, a priori directional 1-tailed t-test ^§^p < 0.05; ^§§^p < 0.01: Ts,*Dyrk1a*^+/+/+^ > Ts,*Dyrk1a*^+/+/Dox-Cre^, a priori directional 1-tailed t-test ^^^p < 0.05; Eu,*Dyrk1a*^+/Dox-Cre^ > Eu,*Dyrk1a*^+*/*+^, post hoc 2-tailed t-test

For P21 mice, DYRK1A protein (Fig. 8C) was overexpressed in the hippocampus and cerebral cortex in Ts,*Dyrk1a*^+/+/+^ as compared to Eu,*Dyrk1a*^+/+^ mice both in females and in males. The functional reduction of *Dyrk1a* copy number at P0 in male Ts,*Dyrk1a^+/+^*^Dox-Cre^ mice significantly reduced DYRK1A expression in the cerebellum at P21 as compared to Ts,*Dyrk1a*^+/+/+^ mice. Unexpectedly, DYRK1A expression was significantly increased in the hippocampus of female Eu,*Dyrk1a*^+/Dox-Cre^ as compared to female Eu,*Dyrk1a*^+/+^ pups. For *Dyrk1a* mRNA on P21 (Fig. 8D), significant overexpression in female mice was evident in the hippocampus and cerebellum of Ts,*Dyrk1a*^+/+/+^ as compared to Eu,*Dyrk1a*^+/+^ littermates, whereas in male mice at P21, mRNA was significantly overexpressed in all three regions in Ts,*Dyrk1a*^+/+/+^ as compared to Eu,*Dyrk1a*^+/+^ littermates.

## DISCUSSION

This is the first report of a DS mouse model genetically engineered to permit a temporally specific, inducible functional reduction of *Dyrk1a* copy number in an otherwise trisomic Ts65Dn mouse model, together with an inducible model of *Dyrk1a* haplo-insufficiency in otherwise euploid mice to test neurobehavioral deficits. The validity of the Ts,*Dyrk1a*^+/+/+^ model was verified by the significant developmental delays (as compared to Eu,*Dyrk1a*^+*/*+^ littermates) in multiple domains assessed in the comprehensive and multi-faceted battery of neurodevelopmental phenotypes. These included reduced postnatal weight (Fig. 1), delayed achievement of forward locomotor competence (Fig. 2) and of multiple physical and sensorimotor milestones (Fig. 3), shifts in developmental regulation of isolation-induced USVs (Fig. 4) and delayed thermoregulatory response to the environmental temperature challenge during isolation (Fig. 5), and the emergence of increased locomotor activity phenotypes in the third postnatal week (Fig. 6).

The functional reduction of one copy of the triplicated *Dyrk1a* gene in the Ts,*Dyrk1a*^+/+/Dox-Cre^ mice, confirmed by sequence-specific PCR after treatment with doxycycline, did not significantly improve the majority of the DS-related neurodevelopmental phenotypes evident in Ts,*Dyrk1a*^+/+/+^ littermates. Two notable exceptions were the significant improvement in the emergence of running in male Ts,*Dyrk1a*^+/+/Dox-Cre^ mice on P19 (Fig. 2), and the reduction of the rate of USV calls during the second postnatal week in Ts,*Dyrk1a*^+/+/Dox-Cre^ mice of both sexes (Fig. 4).

The gene expression studies of *Dyrk1a* mRNA and protein identified significant overexpression in P6 male Ts,*Dyrk1a*^+/+/+^ as compared to Eu,*Dyrk1a*^+/+^ mice in hippocampus, cerebral cortex, and cerebellum; females showed significant overexpression of mRNA in all three regions on P6, but for protein only the cerebral cortex reached significance. For P6 gene expression, the only significant effect due to functional reduction of *Dyrk1a* copy number was in male DYRK1A protein in the cerebellum (Fig. 8 A-B). The gene expression data of P21 mice (Fig. 8 C-D) was assessed in mice that had completed the daily behavioral test battery. Female Ts,*Dyrk1a*^+/+/+^ mice displayed overexpression of protein in hippocampus and cortex (but not cerebellum) and overexpression of mRNA in hippocampus and cerebellum (but not cortex). Male Ts,*Dyrk1a*^+/+/+^ mice also showed overexpression of protein in hippocampus and cortex cerebellum (but not in cerebellum) and overexpression of mRNA in all three regions. The only significant reduction in gene expression on P21 found in Ts,*Dyrk1a*^+/+/Dox-Cre^ as compared to Ts,*Dyrk1a*^+/+/+^ mice was in cerebellar DYRK1A protein in males, similar to results on P6.

Previous reports have described the effects of normalizing *Dyrk1a* from conception in a trisomic system and have detailed improvements in several brain and behavior phenotypes in this DS mouse model [21,57]. Additional experiments have administered pharmacological DYRK1A inhibitors both in humans and mouse models but have produced conflicting results, limited by uncertain target specificity, variations in the age and sex of the test subjects, and measures of success that are either undefined or lack specificity [52,58,59]. The genetic approach of the current study using a temporally controlled excision of one *Dyrk1a* allele to reduce *Dyrk1a* gene copy number can provide a perspective of the likely range of potential improvements in phenotypic outcomes that may result from targeting trisomic DYRK1A, including effects on molecular, growth, and sensorimotor measures that may comprise measures of effects of intervention treatment administered on the day of birth in Ts65Dn neonatal pups. Yet, the current study shows that reducing *Dyrk1a* gene copy number from 3 to 2 on the day of birth elicits only limited rescue effects in many of the neurodevelopmental effects assessed and had limited effects on *Dyrk1a* gene expression as well. These outcomes indicate that the normalization of *Dyrk1a* copy number at birth was insufficient to produce a broad improvement to abnormal phenotypes that emerge during the early postnatal period of development in this Ts65Dn model. It now becomes essential to gain better understanding of the spatiotemporal regulation of *Dyrk1a* and how its dysregulation affects interactions with other developmentally regulated gene networks to produce the neurodevelopmental delays in mouse models of DS.

Infants with DS are typically born underweight [26]. Similarly, low birth weight and lower overall weight in trisomic mouse pups during the postnatal period have been documented in previous studies and results from this study support those previous findings [17,28,60,61]. Both male and female Ts,*Dyrk1a*^+/+/+^ pups had significantly lower birth weight and presented slower growth as compared to controls of the same age and sex throughout the first three weeks. Ts,*Dyrk1a*^+/+/+^ pups gained less weight daily, especially in the first and third postnatal weeks (Fig. 1). Normalizing *Dyrk1a* gene copy number on the day of birth did not rescue the lower weights of trisomic pups, as there were no significant differences in the weights of Ts,*Dyrk1a*^+/+/+^ and Ts,*Dyrk1a*^+/+/Dox-Cre^ pups from P3 to P21.

Delayed physical development is a hallmark characteristic of DS, signs of which are recapitulated in the results of this study. Both male and female Ts,*Dyrk1a*^+/+/+^ compared to Eu,*Dyrk1a*^+*/*+^ mice were delayed by about two days in average day to achieve eye opening, in contrast to previous experiments in which no differences in eye opening were found either in non sex-stratified Ts65Dn pups (line 001924) [25] or in male and female Ts65Dn pups from line 005252 [32]. The delay in eye opening in the current study may have contributed to differences between euploid and trisomic groups on tests occurring between P13-P15, such as the P13 locomotor activity test in which Ts,*Dyrk1a*^+/+/+^ groups were less active and spent more time in the center of the chamber than Eu,*Dyrk1a*^+*/*+^ groups.

Children with DS also have delays in motor skills [9]. Co-activation of brain regions such as the cerebellum and dorsolateral prefrontal cortex may explain the relationship between motor and cognitive development [62]. Male and female Ts,*Dyrk1a*^+/+/+^ as compared to euploid mice exhibited significant developmental delays in each measure of motor development, including pivot, crawl, transition, walk, and run (ambulation milestones), locomotor activity (P10-P16, before hyperactivity at P19), and motility at P11 on the home preference test. In some cases, this delay in Ts,*Dyrk1a*^+/+/+^ mice was profound, as illustrated by the trisomic delay of 5 days in the cliff aversion achievement. The delay of USV peak call number by ∼3 days in Ts,*Dyrk1a*^+/+/+^ as compared to Eu,*Dyrk1a*^+*/*+^ mice may also have a motor component to it, as diaphragm and laryngeal motor control is needed produce these calls. Temporally specific functional reduction of *Dyrk1a* copy number in trisomic mice had only limited effects on correcting this pervasive motor delay. The difference in weight between Ts,*Dyrk1a*^+/+/+^ and Eu,*Dyrk1a*^+*/*+^ mice throughout testing may influence the motor ability of trisomic pups because they lack the strength or development needed to achieve these milestones. Alterations in CNS, rather than musculoskeletal development, may have a significant impact on these delays in motor milestone achievement in trisomic mice.

The delays in motor development in both male and female Ts,*Dyrk1a*^+/+/+^ as compared to Eu,*Dyrk1a*^+*/*+^ mice may implicate functional consequences of cerebellar dysmorphology in Ts65Dn mice [20]. Because cliff aversion involves movements that are less practiced in early development that require strength, coordination, and labyrinthine reflexes, significant weight disparities and delays in sensory and motor development in trisomic pups may particularly contribute to the large delays in cliff aversion achievement [63]. Tactile stimulation tests measure sensory input and coordinated motor reactions [64], and adult Ts65Dn mice are known to have motor coordination impairments [28]. Delays in tactile stimulation achievement, among other motor delays, in both male and female pups, suggest that motor coordination impairment seen in adult Ts65Dn mice may have origins in early development, as suggested by findings in children with DS [9]. The contact righting and negative geotaxis tests assess vestibulo-motor function and development of the pups [63,65]. Absence of vestibular information in early life is linked to reduced cognitive performance, possibly due to the importance of vestibular information input to the hippocampus [66]. Dysfunctional vestibular development may contribute to deficits in delayed contact righting, negative geotaxis, and cliff aversion achievement in both male and female Ts,*Dyrk1a^+/+/+^* pups could be early signs that predict later cognitive impairments.

At P14, male Ts,*Dyrk1a*^+/+/+^ mice showed significant deficits in preference for home cues as compared to male Eu,*Dyrk1a*^+/+^ mice, an effect not seen in females. Sex differences in the presence or extent of some milestone delays were also evident. Male Ts,*Dyrk1a*^+/+/+^ pups showed a delay in the achievement of the Preyer reflex (startle response), similar to previous findings [32] and suggestive of impaired sensory system maturation, whereas no significant delays in age of achievement of the Preyer reflex were seen in trisomic females. In contrast, female as compared to male Ts,*Dyrk1a*^+/+/+^ pups exhibited trends for longer delays with larger effect sizes in achievement of sensorimotor milestones for running, contact righting, tactical stimulation, and negative geotaxis achievements. Given the lack of significant differences in body weight between male and female mice in this study, body size would not appear to account for instances of sex differences in delays in age of achievement of several physical and sensorimotor milestones. Some of these results also contrasted with those seen with the Ts65Dn 005252 DS mouse model, in which female trisomic mice achieved most milestones before males [32]. Whether those different outcomes reflect differences between the two Ts65Dn lines would require direct experimental comparisons.

The altered expression of *Dyrk1a* may also relate to the differences in particular behavioral outcomes between males and females. Although the temporally specific reduction of *Dyrk1a* did not significantly affect weight in either males or females, male Ts,*Dyrk1a*^+/+/Dox-Cre^ mice started running sooner than the Ts,*Dyrk1a*^+/+/+^ mice. This suggests that genetic copy number normalization may impart some sex-specific neurodevelopmental improvements that may become more evident as the trisomic mice approach adolescence (or beyond).

Neonatal ultrasonic vocalization (USV) evoked by isolation of the pup from the dam, nest, and littermates is regulated by changes in thermal, tactile, and olfactory cues [67–69], and USVs show quantifiable spectrographic changes over early postnatal development [70].

Communicative functions of neonatal USVs serve essential adaptive roles by eliciting maternal attention, retrieval, and caregiving responses [67]. Isolation-induced USVs of male and female Ts,*Dyrk1a*^+/+/+^ pups on P3 and P6 were fewer, shorter, with a higher principal frequency and less power than USVs of Eu,*Dyrk1a*^+*/*+^ pups, but on P9 and P12 the trisomic mice made more calls than their euploid littermates. This developmental shift in Ts,*Dyrk1a*^+/+/+^ mice from lower rates of isolation-evoked USVs in the first postnatal week to higher rates in the second postnatal week (Fig. 4) replicates the effect originally reported in the first analysis of isolation-induced USVs in male Ts65Dn mice [25], and extends those findings to females. The current study also demonstrates that the developmental shift in USV calls overlaps with a developmental delay in the emergence of thermoregulatory response to isolation at room temperature of the Ts,*Dyrk1a*^+/+/+^ pups, with both sexes showing significantly impaired development of defense of body temperature (relative to Eu,*Dyrk1a*^+*/*+^ pups) on P6, P9, and P12.

Because environmental temperatures and associated thermoregulatory challenges are predominant determinants of pup USV production during this period, the developmental shift in trisomic mice in USV calling profiles and delayed maturation of homeostatic thermoregulatory mechanisms indicate a significant impairment in the development of these important adaptive functions. Notably, the Ts,*Dyrk1a*^+/+/Dox-Cre^ pups called less than Ts,*Dyrk1a*^+/+/+^ pups in the second postnatal week (thus being more similar to Eu,*Dyrk1a*^+*/*+^ pups) without a change in their thermoregulatory response profile. This suggests that *Dyrk1a* copy number normalization may have affected the developmental trajectory of USV communication but not thermoregulatory mechanisms in response to isolation. Notably, *Dyrk1a* copy number reduction in the Ts,*Dyrk1a*^+/+/Dox-Cre^ female pups resulted in USVs with shorter call length and a higher principal frequency than those of Ts,*Dyrk1a*^+/+/+^ female pups, deviating even farther from euploid levels. These acoustic characteristics in female Ts,*Dyrk1a*^+/+/Dox-Cre^ pups may be indicative of a communication change consistent with sensitized isolation-induced USV calling that occurs in higher frequency ranges [71].

Our previous report using experimentally naïve mice found that at P6, both male and female Ts65Dn mice exhibited DYRK1A protein overexpression in the hippocampus, cerebral cortex, and cerebellum, whereas at P21 Ts65Dn male mice showed DYRK1A overexpression in all three regions but Ts65Dn female mice did not show overexpression in any brain region [50]. The current study only partially replicated those outcomes. At P6, male Ts65Dn mice showed the expected significant overexpression of DYRK1A protein in all three regions and female Ts65Dn mice showed significant overexpression only in the cerebral cortex, with non-significant trends for overexpression in the hippocampus and cerebellum. At P21, male Ts,*Dyrk1a*^+/+/+^ as compared to Eu,*Dyrk1a*^+/+^ mice showed the expected significant overexpression of DYRK1A protein in the hippocampus and cerebral cortex (but not the cerebellum). Unlike our previous report, female Ts,*Dyrk1a*^+/+/+^ P21 mice also showed significant overexpression in the hippocampus and cerebral cortex (but not the cerebellum).

In the current study, *Dyrk1a* mRNA expression on P6 was significantly greater in Ts,*Dyrk1a*^+/+/+^ mice as compared to Eu,*Dyrk1a*^+/+^ mice for all three brain regions both in males and in females. This replicated the findings of Hawley et al. (2024) of mRNA overexpression on P6 in the cerebral cortex and cerebellum in trisomic mice of both sexes, but differences in *Dyrk1a* mRNA levels in the hippocampus did not reach significance in the previous study. On P21 in the current study, *Dyrk1a* mRNA was significantly overexpressed in all three regions in male Ts,*Dyrk1a*^+/+/+^ as compared to Eu,*Dyrk1a*^+/+^ mice; in female Ts,*Dyrk1a*^+/+/+^ mice, significant overexpression of *Dyrk1a* mRNA was found in the hippocampus and cerebellum but not the cerebral cortex. The previous study did not assess mRNA at P21.

Differences in the extent of overexpression of DYRK1A protein in trisomic mice in the current study as compared to our previous study [50], particularly on P21, may be due to differences in the developmental experiences of the animals. In this study (starting at P3), all mice assessed for gene expression on P21 were handled and underwent a series of behavioral tests across days that included varying degrees of potential stress (e.g., handling; separation from dams/littermates; exposure to novel conditions; temperature challenges). The previous study simply harvested brain tissue from pups that were housed with dams and littermates undisturbed except for routine animal care until they were taken at P6 or P21 [50]. The potential for experience-dependent reduction of expression and kinase activity of DYRK1A has been demonstrated in adult transgenic Tg*Dyrk1A* mice (that overexpress DYRK1A) given environmental enrichment [72] and in cystathionine beta synthase-deficient adult mice (that overexpress DYRK1A) given environmental enrichment + voluntary exercise [73]. The extent to which the DYRK1A expression or kinase activity in DS mouse models may be modulated by differential experience (including developmental stressors as in this study) over the first three postnatal weeks is not known.

Although mRNA levels are typically considered an indicator of gene expression that may correlate with protein expression in the same cell, mismatches often occur that may be due to differences in post-translational processing of mRNA, epigenetic regulation, or variations in protein half-life that may yield spatiotemporal complexities that uncouple direct correlation [74,75]. It has been previously shown in Ts65Dn mice that DYRK1A protein and mRNA overexpression may not show typical correlations [50]. For example, in that study P6 hippocampal tissue in both males and females showed significant DYRK1A overexpression, but neither sex showed significant mRNA overexpression in hippocampal tissue at P6. Additionally, DYRK1A protein overexpression was found in the P0 forebrain and cerebellum only in female Ts65Dn, but both male and female tissues showed significant *Dyrk1a* transcript overexpression

The expected reduction of DYRK1A expression after neonatal functional reduction of one *Dyrk1a* allele in otherwise trisomic mice was only confirmed in the cerebellum of male Ts,*Dyrk1a*^+/+/Dox-Cre^ mice as compared to Ts,*Dyrk1a*^+/+/+^ mice, for which significant reductions in DYRK1A were evident on both P6 and P21. No significant reductions were found in mRNA expression in male Ts,*Dyrk1a*^+/+/Dox-Cre^ mice at either age. Moreover, female Ts,*Dyrk1a*^+/+/Dox-Cre^ mice showed no significant reductions in protein or mRNA in any brain region at either age. Interestingly, our previous analysis of P6 Ts65Dn mice with a constitutive *Dyrk1a* copy number reduction from conception found significant reductions of DYRK1A in Ts,*Dyrk1a*^+/+/-^ mice as compared to Ts,*Dyrk1a*^+/+/+^ in both males and females [50]. Several important implications emerge from our findings that significant reductions of DYRK1A levels following *Dyrk1a* copy number reduction were limited to just the cerebellum in males. First, developmental regulation of DYRK1A expression may have strong links to cerebellar development that may differ between males and females. Second, the mechanistic details of and dynamics of developmental regulation of *Dyrk1a* expression in DS mouse models during the early postnatal period need to be better understood.

In typical neurodevelopment, *Dyrk1a* has been established as an essential component for adaptive development and survival in its role of transcriptional regulation of genes responsible for cell growth and differentiation [41,46,76], and its triplication has been implicated in having a major impact on aberrant brain and behavioral development in Ts21. Nevertheless, characterization of *Dyrk1a*’s influence in early perinatal brain development has been underexplored in DS neurobiology research, and studies continue to rely on assumptions regarding where, when, and by how much its expression is dysregulated in DS mouse models carrying an extra copy of this gene. Optimal developmental regulation of DYRK1A highlights its pivotal role in neurodevelopment; insufficient inhibition of trisomic *Dyrk1a* may yield no rescue effect, whereas excessive inhibition could yield neurodevelopmental deviations further from typical outcomes. *Dyrk1a* null mice die during gestation and *Dyrk1a* haploinsufficient mice from birth show evidence of decreased viability and growth, smaller brain size and abnormal brain cell counts, and reduced memory and learning capacity [76]. Non-trisomic, transgenic mice overexpressing *Dyrk1a* exhibit delayed rostro-caudal maturation, altered motor skill acquisition and hyperactivity and significant impairment in spatial learning and cognitive flexibility [47]. In humans, a mutation in one *DYRK1A* allele produces a disease known as Intellectual Developmental Disorder, Autosomal Dominant 7 (MRD7, OMIM:614104) or DYRK1A syndrome, which manifests with microcephaly, intellectual disability, speech delay, and facial dysmorphisms, similar to typical characteristics seen in individuals with DS [77,78]. For optimal efficacy and safety, correcting abnormal gene expression during sensitive periods of neurodevelopment must be tailored to the specific age, sex, and phenotype and aimed to affect only the targeted gene(s) or protein(s) in question [52].

*DYRK1A* has arisen as a candidate for intervention due to numerous studies confirming its implications in measures of cognitive development [42,79–81] and by the finding that increasing only *Dyrk1a* copy number in mice results in motor and learning disabilities [47,82]. Overexpression of DYRK1A in embryonic mice inhibits proliferation and induces premature neuronal differentiation [41] and dysregulations of *Dyrk1a* copy number cause abnormal cerebellar structure and function, indicated by tasks involving coordination, motor learning, and behavior [37,76]. It is probable that *DYRK1A* is one of these select genes, however it is improbable that its overexpression is the only gene critical to reach the threshold of all DS phenotypic manifestations; normalizing *Dyrk1a* copy number did not significantly improve multiple characteristic developmental delays. Triplicated *Dyrk1a* contributes to trisomic phenotypes, but the extent and method of its involvement is not independent of other triplicated genes in Ts21.

Genetic reduction of one copy of triplicated *Dyrk1a* at birth in this DS mouse model yielded only limited support that normalizing *Dyrk1a* copy number produced beneficial outcomes in early postnatal stages. By extension, the absence of improvements across the various tasks in this study similarly failed to support the prediction that targeted pharmacological inhibition of DYRK1A would ameliorate neurodevelopmental delays present in models of DS. However, even the straightforward prediction that copy number reduction of *Dyrk1a* would produce significant reductions in expression of mRNA and protein was largely unconfirmed, being evident only in P6 and P21 male cerebellar protein. Given that measures of DYRK1A kinase activity was beyond the scope of this study, any future intervention studies targeting trisomic genes need to consider potential sensitive periods of neurodevelopmental overexpression while being informed regarding the specific age, sex, and intended phenotype. For optimal efficacy, interventions need to be aimed at affecting only the pathways involving the targeted gene(s) or protein(s) in question. A triplicated gene, by itself in a DS mouse model, may not be causally responsible for an unfavorable phenotype. A triplicated gene, combined with hundreds of other triplicated genes in a DS mouse model in a cellular environment designed for feedback molecular controls and transcriptional regulators in diploid organisms, may then overwhelm the cell’s ability to compensate or correct for abnormal transcription or translation of dysregulated expression of that gene. Additionally, this study only looked at early postnatal phenotypes and did not examine effects of the *Dyrk1a* genetic normalization after P21. Future studies should concentrate on the effects of *Dyrk1a* normalization in older mice. Identifying how genetic dysregulations in Ts21—including of the *Dyrk1a* gene—cause abnormal phenotypes is essential to understand the gene-to-gene interactions contributing to DS phenotypes.

## MATERIALS AND METHODS

### Animal models and care

Animals were bred and housed in the secure AAALAC-accredited Science Animal Resource Center in the Indiana University (IU) Indianapolis School of Science in a vivarium maintained in temperature and humidity-controlled rooms on a standard diurnal 12:12 light/dark cycle (lights on at 0700). All animals had free access to Laboratory Rodent Diet (unless otherwise indicated) (Cincinnati Lab Supply, Stock Number 5001), water, and given a molded paper rodent hut and nesting material for environmental enrichment. Prior approval was received for all experiments involving animals by the IU Indianapolis Institutional Animal Care and Use Committee (SC298R and SC338R) and were performed in accordance with the NIH Guide for the Care and Use of Laboratory Animals. All animals were sourced from the Jackson Laboratory (Bar Harbor, Maine) unless otherwise specified.

B6EiC3Sn a/A-Ts(17^16^)65Dn/J (Stock number 001924) (Ts65Dn) [17] females were bred with male B6C3F1 animals (Stock number 100010), producing both wildtype (euploid) and trisomic offspring with a ∼50% C57BL/6 and ∼50% C3H/HeJ advanced intercross genetic background. New male B6C3F1 and female Ts65Dn mice were purchased and introduced to the mouse colony approximately every six months to maintain consistent animal production and reduce genetic drift within the colony.

Male B6.*Dyrk1a^tm1Jdc^* mice containing loxP sites flanking exons 5 and 6 of *Dyrk1a* were obtained from Dr. Jon Crispino and crossed with C3H/HeJ mice (Stock number 000659) to produce B6C3.*Dyrk^1atm1Jdc^*mice (*Dyrk1a^fl/wt^*) [83]. These animals were intercrossed to produce homozygous B6C3.*Dyrk1a^fl/fl^* mice having loxP sites on both *Dyrk1a* alleles with a similar ∼50% C57BL/6 and ∼50% C3H/HeJ genetic background as Ts65Dn mice. Female Ts65Dn mice were bred with male homozygous B6C3.*Dyrk1a^fl/fl^* mice to produce trisomic and euploid pups with one “floxed” *Dyrk1a* allele (Ts65Dn.*Dyrk1a^fl/wt^*).

B6N.FVB(Cg)-Tg(CAG-rtTA3)4288Slowe/J mice (Stock Number 016532) and B6.Cg-Tg(tetO-cre)1Jaw/J mice (Stock Number 006234) were purchased and mated with C3H/HeJ mice, intercrossed, and backcrossed to achieve a genetic background similar to the ∼50% C57BL/6 and ∼50% C3H/HeJ advanced intercross genetic background as Ts65Dn mice. Male mice carrying the gene for tetO-cre and homozygous for the reverse tetracycline-controlled transcriptional activator (B6C3.tetO-rtTA) genes were bred to Ts65Dn.*Dyrk1a^fl/wt^* female mice. These matings produced euploid and trisomic pups, with or without the full complement of genes necessary to induce a functional reduction of one copy of *Dyrk1a*.

Adult breeding pairs were housed together until late pregnancy was suspected (calculated from the date the male was introduced to the female cage and by visual inspection of the dam). The male was removed during the final days of gestation to prevent repeat pregnancy during postpartum estrus. Cages were checked twice daily for new pups and the day of parturition designated as P0. Pups remained in the home cage with the dam unless otherwise stated. Both male and female pups were used in all experiments and full litters of pups were collected at the same time. For inclusion in this study, each litter was required to contain a minimum of 3 pups, including at least one trisomic pup of either sex. Litters were collected for these projects from May 2022 – September 2024. A total of 152 animals were used from 39 litters. Seven animals expired or were euthanized due to illness or injury (3 Eu, *Dyrk1a*^+/Dox-Cre^, 2 Ts, *Dyrk1a*^+/+/+^, 2 Ts, *Dyrk1a*^+/Dox-Cre^) during testing; data from these animals were excluded from analysis.

### Genotyping of animals

All animals were genotyped using MyTaq DNA Polymerase Kit (Bioline) according to the manufacturer’s instructions. All animals were genotyped for trisomy with primers (forward) 5’-GTGGCAAGAGACTCAAATTCAAC-3′ and (reverse) 5’-GGCTTATTATTATCAGGGCATTT-3′ with (forward) 5’-AAAGTCGCTCTGAGTTGTTAT-3’ and (reverse) 5’-GGAGCGGGAGAAATGGATATG-3′ as internal controls [19]. Detection of the loxP sites flanking *Dyrk1a* exons 5 and 6 was performed using primers (forward) 5’-TACCTGGAGAAGAGGGCAAG-3′ and (reverse) 5’-GGCATAACTTGCATACAGTGG-3′ [83] or primers (forward) 5’-ATTACCTGGAGAAGAGGGAAG-3′ and (reverse) 5’-TTCTTATGACTGGAATCGCC-3′. Primers (forward) 5’-CTGCTGTCCATTCCTTATTC-3′ and 5’-TGCCTATCATGTTGTCAAA-3′ and (reverse) 5’-CGAAACTCTGGTTGACATG-3′ were used to genotype for the rtTA gene [84]. Genotyping for the tetO-cre gene was accomplished using primers (forward) 5’-ATTCTCCCACCGTCAGTACG-3′ and (reverse) 5’-CGTTTTCTGAGCATACCTGGA-3′ [85]. A separate set of adult animals containing all genetic components for the functional reduction of *Dyrk1a* had tissues collected both before and after the administration of doxycycline feed to confirm the specificity of the excision mechanism and check for a “leaky flox”. The *Dyrk1a* excision mechanism was confirmed to be functional only in animals containing all genetic components and only after receiving doxycycline feed as previously done [86].

Successful truncation of the *Dyrk1a* allele was confirmed in all Ts,*Dyrk1a*^+/+/Dox-Cre^ and Eu,*Dyrk1a*^+/Dox-Cre^ offspring with primers (forward) 5’-ATTACCTGGAGAAGAGGGCAAG-3′ and (reverse) 5’-CCTAGAGCAGCCCACATTCT-3′ and 5’-CAAACAGCCACTGTGTGAGG-3′ which target the span of exons 3 and 7 (without exons 5 and 6) [53,83].

### Inducible knockdown of *Dyrk1a*

Three genetic components, controlled by the presence or absence of doxycycline, were necessary to initiate the functional reduction of one *Dyrk1a* allele in the animal subjects: the reverse-tetracycline transcription activator (CAG-rtTA—from Jackson Laboratory stock # 016532), Cre recombinase under the control of a tetracycline-responsive promoter element (tetO-cre—from Jackson Laboratory stock # 006234), and a floxed *Dyrk1a* allele inserted to flank exons 5 and 6 (from John Crispino, now Jackson Laboratory stock # 027801) spanning the DNA sequence that codes for the active site of the DYRK1A protein. In mice given doxycycline with all three necessary genetic elements, the combination of the tetO and rtTA activated the Cre to excise the floxed portion of *Dyrk1a*. Mothers with full litters of pups from (Ts65Dn.*Dyrk1a^fl/wt^* x B6C3.tetO-rtTA) matings were randomly assigned either doxycycline feed (ENVIGO, Teklad Custom Diet TD.120769, 998.975g/kg 2018 Teklad Global 18% Protein Rodent Diet, 0.625g/kg Doxycycline hyclate, 0.4g/kg) or Laboratory Rodent Diet 5001 (as described above) starting on the day of birth (postnatal day 0 [P0]) until P6. Dams on both diets ate approximately 5g of feed daily. Those receiving doxycycline feed consumed approximately 2-3mg of doxycycline per day through the infused feed, a portion of which passed to the pups via breastmilk [87]. Litters receiving doxycycline treatment produced male and female offspring with four distinct genotypes, with different numbers of functional copies of *Dyrk1a*: euploid (Eu,*Dyrk1a*^+/+^), euploid with a functional *Dyrk1a* reduction (Eu,*Dyrk1a*^+/Dox-Cre^), Ts65Dn (Ts,*Dyrk1a*^+/+/+^), and Ts65Dn with a functional *Dyrk1a* reduction (Ts,*Dyrk1a*^+/+/Dox-Cre^). Tissues from animals of all four genotypes were taken from the heart, liver, lungs, thymus, muscle, kidney, spleen, and lung and PCR performed to confirm *Dyrk1a* was only truncated under the controlled conditions [86]. Additionally, DNA was isolated from the cerebellum, cerebral cortex, and hippocampus of each of the four genotypes to confirm controlled *Dyrk1a* excision in each brain region. The *Dyrk1a* excision mechanism was confirmed by collecting tissues of matched animals before and after administration of doxycycline feed.

### Data analysis approach

In keeping with the three a priori hypotheses afforded by this genetic approach, measures of most of the phenotypes (described below) were analyzed with three different statistical comparisons, conducted separately for females and males. For the first a priori hypothesis, differences between trisomic (Ts,*Dyrk1a*^+/+/+^) and euploid (Eu,*Dyrk1a*^+/+^) mice were tested with directional, one-tailed t-tests (in the appropriate direction, e.g., later milestone achievement in DS mice). In cases involving repeated measures such as daily body weight, ultrasonic vocalizations (USVs) over days, or locomotor activity over a session, repeated measures ANOVAs were conducted for Ts,*Dyrk1a*^+/+/+^ and Eu,*Dyrk1a*^+/+^ groups when specific directional effects of genotype were expected. For example, for USVs, previous findings indicated that Ts65Dn mice emitted fewer calls on P6 but more calls on P9 than euploid mice, thus predicting a genotype × age interaction. When specific ANOVAs were appropriate for the a priori hypotheses, significant main effects in the predicted direction in the absence of a significant interaction were reported. When significant interactions were present, follow-up simple main effects contrasts were used to identify significant group differences between the two genotypes at each value of the repeated measure.

The second a priori hypothesis tested the potential improvement due to the functional reduction of *Dyrk1a* in otherwise trisomic mice by comparing the Ts,*Dyrk1a*^+/+/Dox-Cre^ and Ts,*Dyrk1a*^+/+/+^ mice. These used a similar directional hypothesis testing approach described above (either directional one-tailed t-tests or two-group repeated ANOVAs) by assessing whether effects evident in Ts,*Dyrk1a*^+/+/+^ mice were significantly diminished in the Ts,*Dyrk1a*^+/+/Dox-Cre^ mice . Likewise, the third a priori hypothesis tested the potential effect of a functional reduction of *Dyrk1a* in otherwise euploid mice (Eu,*Dyrk1a*^+/Dox-Cre^) as compared to euploid littermates (Eu,*Dyrk1a*^+/+^). Given that the direction of effects of haploinsufficiency are less well known and often not previously reported in mouse models, these analyses used two-tailed non-directional t-tests instead of directional t-tests. The specific dependent variables and analyses are identified in the description of each test. Statistical analyses were performed using IBM SPSS for Windows version 29.0.1 or Prism GraphPad, and graphs were created in Prism GraphPad.

### Protein and mRNA isolation and quantification

At P21 (after all behavioral testing was completed), the full litter of pups was separated from the dam, pups were euthanized via isoflurane inhalation and cervical dislocation before the hippocampus (HIP), cerebral cortex (CTX), and cerebellum (CB) were collected from all male and female pups that underwent behavioral testing. Similar brain tissue was collected from P6 animals that had not undergone behavioral testing. Tissues were separated by left and right halves, snap frozen in liquid nitrogen, and stored at -80°C until further processing; left or right halves of each tissue were chosen at random for protein or mRNA isolation.

Protein isolation and Western blot quantification were performed as previously described [50,53]. In short, 20µg of protein was resolved electrophoretically and transferred to a PVDF membrane. After blocking in 5% Blotting Grade Blocker (Bio-Rad), membranes were incubated in 1:500 DYRK1A monoclonal antibody (M01), clone 7D10 (Abnova) followed by Donkey anti-mouse IgG (H + L) Highly Cross-Adsorbed Secondary Antibody, Alexa Fluor™ 790 (ThermoFisher). For P21 animals, protein of the HIP and CTX of six animals, almost exclusively of one sex, were run concurrently on one blot, along with three lanes of 20µg aliquots of a protein homogenate (used in all blots) that combined the HIP, CTX, and CB of 6 male and 6 female P6 euploid mice. The samples from the common homogenate served as a comparison standard for quantifying relative amounts of DYRK1A across all blots. For the CB, 12 animals of a single sex were run concurrently on one blot along with three lanes of the same homogenate control. For P6 animals For the P6 blots, all three brain regions of the same pups were run on the same blot together. The P6 blots were also almost exclusively of one sex. Relative DYRK1A protein expression of each sample was calculated as a relative intensity ratio of the normalized value of each sample (DYRK1A fluorescence / total protein) divided by the mean of the normalized values of the three homogenate standards from the same blot. Within sex and brain region, three a priori directional hypotheses regarding DYRK1A protein levels were tested, Eu,*Dyrk1a*^+/+^ < Ts,*Dyrk1a*^+/+/+^, Ts,*Dyrk1a*^+/+/Dox-Cre^ < Ts,*Dyrk1a*^+/+/+^, and Eu,*Dyrk1a*^+/Dox-Cre^ < Eu,*Dyrk1a*^+/+^, using one-tailed independent t-tests (p < 0.05), corrected for unequal variances when Levene’s test was significant. Note that directional one-tailed tests were used for the a priori hypothesis for euploid groups because the predicted effect of copy number reduction is a reduction in expression.

RNA was isolated from the HIP, CTX, and CB of the same animals used for protein isolation, using the other half of the randomly selected left and right brain tissues as previously described [50]. Briefly, samples were incubated on ice for 30 minutes in TRIzol Reagent (ThermoFisher) before being homogenized by sonification, centrifuged, and the resulting supernatant removed to new tubes. Chloroform was added to separate the aqueous phase, which was transferred to a second set of new tubes. Samples were washed first in 100% ice-cold isopropanol followed by a second wash in 75% ice-cold ethanol. The resulting pellet was air dried for 5 minutes and rehydrated with 30µL nuclease-free water and stored at -80°C. Samples were quantified using a NanoDrop 2000 (Thermo Scientific). RNA samples were diluted to 200-500 ng/μL using RNase/DNase free water for more accurate cDNA conversion. 100ng of each RNA sample was converted to cDNA using TaqMan Reverse Transcription Reagents (ThermoFisher Scientific) according to the manufacturer’s instructions. After conversion, cDNA samples were diluted in sterile MilliQ water in a 1:5 ratio and stored at -20°C.

RT-qPCR was performed using TaqMan Gene Expression Master Mix (ThermoFisher Scientific) with a *Dyrk1a* probe targeting the span between exons 5 and 6 (ThermoFisher Mm00432929_m1) and a probe targeting the 18S ribosomal RNA (ThermoFisher Mm03928990_g1), which was used as a universal standard for ΔΔCt calculations as described [50]. Verification of qPCR efficiency for the *Dyrk1a* and *Rn18s* probes was performed with serial dilution curves using tissue from the hippocampus, cerebral cortex and cerebellum of two Eu^+/+^ and two Ts^+/+/+^ P21 female mice on one plate and of two Eu^+/+^ and two Ts^+/+/+^ P21 male mice on a second plate. For each brain region of each animal, triplicated replicates of undiluted cDNA and four tenfold serial dilutions were amplified for each probe. The standard deviations of replicates over the five dilutions for *Dyrk1a* and *Rn18s*, respectively, averaged 0.136 and 0.126 for females and 0.124 and 0.080, respectively, for males. After amplification, the cycle threshold versus log cDNA dilution was plotted separately for females and males to determine the log-linear regression for each probe for each of the three brain regions of the two euploid and two trisomic mice of each sex. The slope of each regression line of those plots was used to determine PCR efficiency. For females (Supplemental Fig. 1 and Supplemental Table 1), the mean (± SEM) efficiency was 92.0% (± 0.9) for *Dyrk1a* and 97.2% (± 0.7) for *Rn18s*; for males (Supplemental Fig. 2 and Supplemental Table 2), the mean (± SEM) efficiency was 94.2% (± 0.8) for *Dyrk1a* and 96.6% (± 0.67) for *Rn18s*. The efficiency was verified to be above 90% and the differences between the reference and target probe were within ten percent of each other.

The relative expression of *Dyrk1a* was determined as in our previous report [50] using the 2^-ΔCT^ comparative CT method [88]. For each sample, 2^-ΔCT^ was first calculated [2^-(CT*Dyrk1a*−CT*Rn18s*)^] and that value served as the numerator of the relative ratio calculation for the sample with the denominator being the mean of the samples of the Eu,*Dyrk1a^+/+^* mice of the same brain region from the same plate. These relative expression ratios for each sample yield a measure of *Dyrk1a* expression as a fold change relative to the euploid control mean (of 1.0) to quantify the relative fold increase (values >1.0) or decrease (values <1.0) in expression. For statistical analysis of the qPCR data within sex and brain region, the three previously described a priori directional hypotheses were tested: Eu,*Dyrk1a*^+/+^ < Ts,*Dyrk1a*^+/+/+^, Ts,*Dyrk1a*^+/+/Dox-Cre^ < Ts,*Dyrk1a*^+/+/+^, and Eu,*Dyrk1a*^+/Dox-Cre^ < Eu,*Dyrk1a*^+/+^, using one-tailed independent t-tests (p < 0.05), corrected for unequal variances when Levene’s test was significant.

### Physical, Sensorimotor, and Neurobehavior Development Assessments

To quantify differences between male and female Ts65Dn and euploid mice and evaluate the influence of functional *Dyrk1a* gene copy number on growth and behavior in a trisomic system, a battery of physical, sensorimotor, and neurobehavioral assessments was used to test offspring from P3 through P21 using previously published and adapted protocols [63,89,90] (see Supplemental Fig. 3).

### Physical Development

Animals were maintained on a standard 12:12 light/dark cycle (lights on at 7:00am) and all testing was performed in a room with overhead fluorescent lighting (136 lux). Testing was initiated no earlier than 7:00am and completed no later than 11:00am each day. Pups were weighed daily to the nearest hundredth gram from P3 to P21. The home cage containing the dam and pups was covered with a dark cloth and brought from the vivarium to a cleaned testing room and the animals were allowed to acclimate ∼20 minutes prior to testing. Pups were tested one at a time on a 33.5cm × 26.3cm flexible plastic mat with a 1cm × 1cm grid which was maintained at 36°C using a water circulating heating pad. When testing was completed on P3, each pup received an identification tattoo on one or more of its footpads before being returned to the nest.

Physical growth was evaluated and milestones recorded as achieved on the postnatal day when the developmental milestone met preset criteria. Pinnae detachment was recorded as the postnatal day when the tips of both ears were detached from the sides of the head [90]; in cases where detachment occurred before testing began, the day of achievement was recorded as P3. Upper and lower incisor eruption was recorded as the postnatal day when both incisors had erupted from the gingiva as a set, and eye opening recorded as the postnatal day when both eyes were fully open [91]. Contact righting [65] was used in place of a typical surface righting response, as 96% of all tested pups had achieved the surface righting response milestone before P3 when testing began. Contact righting was performed by placing a pup nose first into a 50mL conical tube lying horizontally on the testing surface. The pup was allowed to acclimate for 5sec before the tube was quickly rolled until all four limbs of the pup were facing up and the pup was fully on its back inside the tube. The milestone was marked as achieved on the second of two days in a row when the pup successfully rolled over onto its feet within 5sec, three times in a row. Tactile stimulation was performed by placing a 1mm circular sticker on the end of the nose and counting the number of seconds required for the pup to successfully remove the sticker [64]. The milestone was marked achieved on the second of two consecutive days when the pup removed the sticker within 10sec of being released. Cliff aversion was measured by the response to being placed with its nose and front paws over the edge of a cardboard box and marked as achieved on the second of two days in a row that the pup moved its body away from the edge of the box within 10sec, three times in a row [91]. Negative geotaxis was performed by placing the pup nose-down on a metal screen platform set at a 45-degree angle. The pup was required to turn its body a minimum of 90-degrees towards the top of the platform within 20sec, three times in a row for two days in a row [91]. The Preyer response was tested by clicking a metal noisemaker ∼4.5cm above the head and the test was repeated three times with an interval of approximately 15 seconds between each click. The day of achievement was recorded as the second of two consecutive days when all three clicks elicited an ear or body twitch response [90].

Daily weight over time and percent body weight changes each day were analyzed separately for males and females using repeated measures analyses of variance (RM-ANOVA) that tested each of the three a priori hypotheses, with postnatal day as the repeated measure. Body weight gained by day was calculated as a percentage difference from the previous day’s weight [((Weight on postnatal day (X)) – (Weight on postnatal day (X-1))) / (Weight on postnatal day (X-1))]*100. For milestone achievement, the first two a priori hypotheses (Eu,*Dyrk1a*^+/+^ achieves milestones before Ts,*Dyrk1a*^+/+/+^; Ts,*Dyrk1a*^+/+/Dox-Cre^ achieves milestones before Ts,*Dyrk1a*^+/+/+^) were tested using directional one-tailed independent t-tests (p < 0.05), corrected for unequal variances when Levene’s test was significant. The third a priori hypothesis (Eu,*Dyrk1a*^+/Dox-Cre^ will differ from Eu,*Dyrk1a*^+/+^) used two-tailed independent t-tests

### Ambulation

Trained animal technicians blind to genotype status tested pups daily, one at a time, on a 33.5cm × 26.3cm flexible plastic mat with a 1cm × 1cm grid warmed by a water circulating heating pad set to 36°C. Unstimulated movement was categorized each day. Movement was classified into five categories based on previously published protocols [63,90] using the following criteria: Pivoting was achieved when the pup used one or both front limbs to push against the surface, achieving at least a 90-degree rotation in either direction. Crawling was classified as the pup moving in a straight line, without directional intent, with its belly, full footpad, and leg to the first joint flat on the surface as it moved. The criteria for achieving the transition phase included movement in a straight line with observable directionality, the belly held off the surface, and the tail held low. Additionally, the toes and footpads, but not the first joint of the leg, were in contact with the surface with each step. Walking was recorded when the pup moved in a straight line with directional intent, tail held high, with only the toes of the feet in contact with the surface and only one foot leaving the surface at a time during each stride. The pup was classified as running when it lifted more than one foot off the ground during each stride. For these measures of forward movement, the first two a priori directional hypotheses (Eu,*Dyrk1a*^+/+^ achieves milestones before Ts,*Dyrk1a*^+/+/+^; Ts,*Dyrk1a*^+/+/Dox-Cre^ achieves milestones before Ts,*Dyrk1a*^+/+/+^) were analyzed using one-tailed independent t-tests (p < 0.05), corrected for unequal variances when Levene’s test was significant. The third hypothesis (Eu,*Dyrk1a*^+/Dox-Cre^ differ from Eu,*Dyrk1a*^+/+^groups) was tested using two-tailed independent t-tests.

### Ultrasonic Vocalization (USV)

#### Isolation-induced USV generation during first two postnatal weeks

On P3, P6, P9, and P12 a home cage containing a dam and a full litter of pups was covered with a dark cloth and brought from the vivarium to a lighted testing room within the animal facility. The cage was placed on a 36°C water-circulating heating pad, and the dam and litter were allowed to acclimate for 20 min while the testing chambers were cleaned with 70% ethanol and recording equipment set up and calibrated. A single pup was removed from the home nest, identified, and its body temperature immediately measured with 0.1°C resolution using a digital infrared thermometer placed directly on its belly. The pup was then placed feet-down in a 500mL plastic beaker in a dark, sound-attenuating chamber insulated with acoustic charcoal foam (BRS/LVE, Laurel, MD), and the audio recording started as soon as the door was latched. The interval between pup removal from the nest and the start of the USV recording was less than 10sec. USVs were recorded for 5min on an Apple iPhone 5 (Apple Inc., Cupertino, CA) using an Echo Meter Touch Ultrasonic Microphone Module and the accompanying Echo Meter Touch application (Wildlife Acoustics, Maynard, MA). At the end of the 5min session, the pup was removed from the chamber and body temperature measured again before being returned to the home cage with its dam and siblings.

Prior to analysis, recordings from six euploid pups at each age (3, 6, 9, and 12 days and taken from litters outside the testing sets) were used to construct a neural network source file. Calls from these 24 audio files were converted and analyzed using DeepSqueak [92], a deep learning software created to analyze USVs in rodents using vision algorithms to convert audio recordings into 2D graphic representations (sonograms) (Supplemental Fig. 4). A trained technician identified the vocalizations within each recording using DeepSqueak software, outlining the parameters for each call that would be used to analyze and measure all subsequent sessions. Five parameters were identified for evaluation within each 5min recording session: 1) the number of calls emitted, 2) the average call length (duration), 3) the average peak frequency (kHz) (the maximum intensity frequency), 4) the average principal frequency (kHz) (the most dominant frequency) and 5) the average power (dB/kHz). The data taken from these 24 representative sessions were used to perform a deep learning session to train the source file against which all USV recordings were analyzed. This trained source file allowed for the direct comparison of euploid and non-euploid call profiles within the same testing conditions, age ranges, and genetic background of all subjects. All USV recordings for each test subject were run through DeepSqueak against the trained source file and the resulting data were exported into Microsoft Excel files for curation and analysis. Results were analyzed using one-tailed independent t-tests (p < 0.05), corrected for unequal variances when Levene’s test was significant, for the first two a priori directional hypotheses (Ts,*Dyrk1a*^+/+/+^ is delayed compared to Eu,*Dyrk1a*^+/+^ and Ts,*Dyrk1a*^+/+/+^ is delayed compared to Ts,*Dyrk1a*^+/+/Dox-Cre^) and using two-tailed independent t-tests for the third a priori hypothesis (Eu,*Dyrk1a*^+/Dox-Cre^ ≠ Eu,*Dyrk1a*^+/+^.

#### Body Temperature Effects During USV Testing

In the first two postnatal weeks, removal of rodent pups from the nest and isolating them at room temperature provokes a thermoregulatory challenge, and loss of body temperature is a major component in evoking isolation-induced USVs [69]. In turn, USV generation is associated with heat production in the pup [93]. Given the important links between body temperature and isolation-induced USV production during this developmental period, the pre- and post-USV test body temperature measures obtained for each session assessed the extent to which DS model mice showed developmental deficits in thermoregulatory responses to isolation at room temperature during the USV test. The extent to which trisomic mice display thermoregulatory deficits associated with age-dependent differences in tests of isolation-induced USV production has not been previously reported. For each of the three a priori hypotheses, the pre- and post-test body temperature data were analyzed with RM-ANOVAs (with day and pre/posttest as repeated measures) followed by Sidek post hoc comparisons.

### Locomotor Activity

On P10, P13, P16, and P19 pups underwent a 10min locomotor activity assessment as previously described [53]. Cages with both dam and pups were brought from the vivarium to a testing room and allowed to acclimate in ambient light for ∼20min while the testing chambers (Versamax Animal Activity Monitors, Accuscan Instruments, Columbus, OH) were cleaned with 70% ethanol and recording equipment prepared and calibrated. Testing chambers consisted of an open-top acrylic chamber measuring 40cm × 40cm × 40cm situated inside a 53cm × 58cm × 43cm closed sound-attenuated box. Chambers were equipped with low, indirect house light (36-41 lux) and a ventilation fan provided fresh air and white noise during testing. The test was started simultaneously with the pup being placed on all four feet in the center of the chamber and the chamber door being closed. Distance traveled was measured as the pup moved within the chamber and interrupted the intersection of photocell beams evenly spaced along all 4 walls of the activity chamber. At the end of each session the pup was returned to the home cage with its dam and siblings, and the testing chamber was disassembled and cleaned with 70% ethanol between subjects. Data were recorded in 1-minute time bins and translated using the Versamax Animal Activity software, which converted the data to a Microsoft Excel document for curation and analysis. Measures recorded included total distance traveled (cm) and the percentage of time the subject spent in the center of the chamber (> 8cm from the walls). Sexes were analyzed separately using RM-ANOVA to measure activity levels at each postnatal age, with time bin (minute) as the repeated measure and genotype (Eu,*Dyrk1a*^+/+^ and Ts,*Dyrk1a*^+/+/+^ , Ts,*Dyrk1a*^+/+/Dox-Cre^ and Ts,*Dyrk1a*^+/+/+^, or Eu,*Dyrk1a*^+/+^ and Eu,*Dyrk1a*^+/Dox-Cre^) as a between-group factor to evaluate significant main effects or interactions of genotype. Significant interactive effects were followed up with simple main effects contrasts to assess differences between genotypes at each time point.

### Home Shavings Preference Test

A version of a home shavings preference test previously described was utilized [53]. On P11 and P14, the cage with both dam and pups was brought from the vivarium to the testing room and allowed to acclimate to the room under ambient light for ∼20min while the testing chamber was cleaned with 70% ethanol and prepared for testing. The chamber consisted of a standard clean mouse cage (28.9 cm long x 16.3 cm wide x 12.2 cm tall) containing two small plastic petri dishes (7 cm diameter, 2 cm tall) in two corners at one end of the cage, with one dish containing clean bedding and the other containing soiled bedding from the home cage, collected the day of testing. A digital video recording of each homing test was recorded with an overhead web camera with a complete view of the testing chamber. Once the recording was started, the pup was placed in the end of the cage opposite the cups and facing them. The recording of the trial then continued uninterrupted for 60 seconds, the recording was stopped, and the pup was returned to its home cage. The testing chamber and cups were cleaned with 70% ethanol, wiped dry, and replenished with the appropriate shavings for the next animal; the location of each cup was switched between each pup.

Two observers blind to ploidy and sex scored each video using the Observer XT version 15 software (Noldus, Wageningen, Netherlands). Three zones were designated that corresponded to the “home zone” (dish containing bedding from home cage), the “clean zone” (dish containing fresh bedding), and the “cage zone” (the rest of the testing cage). Entries into each zone by the pup were scored by keystrokes by the observer, starting when the pup was placed in the start area at the end of the testing chamber farthest from the cups. Total duration and mean duration in a zone, number of entries into each zone, and latency to reach each zone were calculated by the software. The average scores from the two observers were used for data analysis. As testing proceeded, it was recognized that many pups on P11 failed to traverse the testing cage to reach either cup zone. To quantify this apparent reduced mobility, the recordings from P11 and P14 pups were also scored categorically for maximal traversal distance (failed to cross the midline, crossed midline but did not reach cups, entered at least one cup, entered both cups).

Home bedding preference ratio was calculated by (time in home zone)/(time in home zone + time in clean zone). This preference ratio was analyzed using one-tailed independent t-tests (p < 0.05), corrected for unequal variances when Levene’s test was significant, to analyze two a priori directional hypotheses (Ts,*Dyrk1a*^+/+/+^ is delayed compared to Eu,*Dyrk1a*^+/+^ and Ts,*Dyrk1a*^+/+/+^ is delayed compared to Ts,*Dyrk1a*^+/+/Dox-Cre^), and two-tailed independent t-tests were used to analyze Eu,*Dyrk1a*^+/Dox-Cre^ and Eu,*Dyrk1a*^+/+^ comparisons. One-sample t-tests were used to assess whether each group’s average home preference score was significantly different from chance (50%). For the traversal measures, the comparisons of group differences reflecting the three a priori hypotheses were analyzed with non-parametric Fisher’s Exact test.

## CRediT Authorship Contribution Statement

A. D. and L.H. contributed equally to this work and are co-first authors

Alyssa Duerst: Formal analysis, Investigation, Visualization, Writing – original draft preparation, Project administration

Laura Hawley: Conceptualization, Formal analysis, Investigation, Visualization, Writing – original draft preparation, Project administration

Linnea Johnson: Investigation, Formal analysis, Writing – review and editing

Drew Folz: Investigation, Formal analysis, Writing – review and editing

Zaina Obeid: Investigation, Formal analysis, Writing – review and editing

Laura Snellenberger: Investigation, Formal analysis, Writing – review and editing

Charles R. Goodlett: Conceptualization, Resources, Writing – original draft preparation, Funding acquisition, Supervision, Project administration

Randall J. Roper: Conceptualization, Resources, Writing – original draft preparation, Funding acquisition, Supervision, Project administration

## Data Availability

Data have been uploaded to Dryad: http://datadryad.org/share/LINK_NOT_FOR_PUBLICATION/HeQCAZewx5mesochgnRd9Uyv-rXB2_9ppocbH4fZ6f8

