## Supplemental Tables and Figures for "Sex-specific developmental phenotypes and their response to neonatal *Dyrk1a* reduction in the Ts65Dn Down syndrome mouse model"

**SUPPLEMENTAL MATERIAL**

**[TABLES]**

| **Supplemental Table 1: qPCR Efficiency Validation for Females** | | | | | | | | |
| --- | --- | --- | --- | --- | --- | --- | --- | --- |
| **Animal#** | **Sex** | **Genotype** | **Region** | **Gene** | **Slope** | **Y intercept** | **R^2^** | **Efficiency (%)** |
| 990 | Female | Eu | HIP | *Dyrk1a* | -3.632 | 24.17 | 0.9966 | 88.5 |
| 1266 | Female | Eu | HIP | *Dyrk1a* | -3.420 | 24.25 | 0.9965 | 96.1 |
| 1081 | Female | Ts | HIP | *Dyrk1a* | -3.514 | 24.02 | 0.9980 | 92.6 |
| 1221 | Female | Ts | HIP | *Dyrk1a* | -3.609 | 23.84 | 0.9971 | 89.3 |
| 990 | Female | Eu | CTX | *Dyrk1a* | -3.697 | 24.58 | 0.9982 | 86.4 |
| 1266 | Female | Eu | CTX | *Dyrk1a* | -3.533 | 23.29 | 0.9989 | 91.9 |
| 1081 | Female | Ts | CTX | *Dyrk1a* | -3.471 | 24.29 | 0.9991 | 94.1 |
| 1221 | Female | Ts | CTX | *Dyrk1a* | -3.450 | 24.42 | 0.9982 | 94.9 |
| 990 | Female | Eu | CB | *Dyrk1a* | -3.377 | 25.46 | 0.9879 | 97.8 |
| 1266 | Female | Eu | CB | *Dyrk1a* | -3.541 | 23.78 | 0.9981 | 91.6 |
| 1081 | Female | Ts | CB | *Dyrk1a* | -3.506 | 24.47 | 0.9993 | 92.9 |
| 1221 | Female | Ts | CB | *Dyrk1a* | -3.630 | 23.02 | 0.9973 | 88.6 |
| **Mean** |  |  |  |  | -3.532 | 24.13 | 0.9971 | 92.0 |
| **SEM** |  |  |  |  | 0.026 | 0.17 | 0.0008 | 0.9 |
| 990 | Female | Eu | HIP | *Rn18s* | -3.329 | 10.14 | 0.9993 | 99.7 |
| 1266 | Female | Eu | HIP | *Rn18s* | -3.300 | 9.92 | 0.9987 | 100.9 |
| 1081 | Female | Ts | HIP | *Rn18s* | -3.381 | 10.53 | 0.9969 | 97.6 |
| 1221 | Female | Ts | HIP | *Rn18s* | -3.330 | 10.22 | 0.9994 | 99.7 |
| 990 | Female | Eu | CTX | *Rn18s* | -3.467 | 9.71 | 0.9990 | 94.3 |
| 1266 | Female | Eu | CTX | *Rn18s* | -3.452 | 8.20 | 0.9989 | 94.8 |
| 1081 | Female | Ts | CTX | *Rn18s* | -3.399 | 9.49 | 0.9990 | 96.9 |
| 1221 | Female | Ts | CTX | *Rn18s* | -3.307 | 10.18 | 0.9994 | 100.6 |
| 990 | Female | Eu | CB | *Rn18s* | -3.449 | 10.22 | 0.9972 | 95.0 |
| 1266 | Female | Eu | CB | *Rn18s* | -3.435 | 9.02 | 0.9979 | 95.5 |
| 1081 | Female | Ts | CB | *Rn18s* | -3.409 | 10.65 | 0.9983 | 96.5 |
| 1221 | Female | Ts | CB | *Rn18s* | -3.439 | 9.86 | 0.9991 | 95.3 |
| **Mean** |  |  |  |  | -3.391 | 9.85 | 0.9986 | 97.2 |
| **SEM** |  |  |  |  | 0.017 | 0.19 | 0.0002 | 0.7 |

| **Supplemental Table 2: qPCR Efficiency Validation for Males** | | | | | | | | |
| --- | --- | --- | --- | --- | --- | --- | --- | --- |
| **Animal#** | **Sex** | **Genotype** | **Region** | **Gene** | **Slope** | **Y intercept** | **R^2^** | **Efficiency (%)** |
| 857 | Male | Eu | HIP | *Dyrk1a* | -3.450 | 24.82 | 0.9978 | 94.9 |
| 1244 | Male | Eu | HIP | *Dyrk1a* | -3.491 | 23.77 | 0.9983 | 93.4 |
| 861 | Male | Ts | HIP | *Dyrk1a* | -3.419 | 24.24 | 0.9987 | 96.1 |
| 1131 | Male | Ts | HIP | *Dyrk1a* | -3.464 | 23.92 | 0.9985 | 94.4 |
| 857 | Male | Eu | CTX | *Dyrk1a* | -3.323 | 24.79 | 0.9911 | 99.9 |
| 1244 | Male | Eu | CTX | *Dyrk1a* | -3.468 | 24.01 | 0.9988 | 94.2 |
| 861 | Male | Ts | CTX | *Dyrk1a* | -3.515 | 24.15 | 0.9989 | 92.5 |
| 1131 | Male | Ts | CTX | *Dyrk1a* | -3.471 | 24.29 | 0.9991 | 94.1 |
| 857 | Male | Eu | CB | *Dyrk1a* | -3.640 | 23.46 | 0.9935 | 88.2 |
| 1244 | Male | Eu | CB | *Dyrk1a* | -3.416 | 23.58 | 0.9994 | 96.2 |
| 861 | Male | Ts | CB | *Dyrk1a* | -3.495 | 24.07 | 0.9987 | 93.3 |
| 1131 | Male | Ts | CB | *Dyrk1a* | -3.510 | 23.65 | 0.9974 | 92.7 |
| **Mean** |  |  |  |  | -3.472 | 24.06 | 0.9975 | 94.2 |
| **SEM** |  |  |  |  | 0.021 | 0.12 | 0.0007 | 0.8 |
| 857 | Male | Eu | HIP | *Rn18s* | -3.512 | 11.04 | 0.9986 | 92.6 |
| 1244 | Male | Eu | HIP | *Rn18s* | -3.457 | 9.60 | 0.9996 | 94.7 |
| 861 | Male | Ts | HIP | *Rn18s* | -3.294 | 10.07 | 0.9998 | 101.2 |
| 1131 | Male | Ts | HIP | *Rn18s* | -3.368 | 10.23 | 0.9998 | 98.1 |
| 857 | Male | Eu | CTX | *Rn18s* | -3.445 | 10.10 | 0.9999 | 95.1 |
| 1244 | Male | Eu | CTX | *Rn18s* | -3.386 | 9.52 | 0.9997 | 97.4 |
| 861 | Male | Ts | CTX | *Rn18s* | -3.386 | 9.85 | 0.9992 | 97.4 |
| 1131 | Male | Ts | CTX | *Rn18s* | -3.399 | 9.49 | 0.9990 | 96.9 |
| 857 | Male | Eu | CB | *Rn18s* | -3.416 | 10.04 | 0.9998 | 96.2 |
| 1244 | Male | Eu | CB | *Rn18s* | -3.341 | 10.99 | 0.9995 | 99.2 |
| 861 | Male | Ts | CB | *Rn18s* | -3.447 | 9.32 | 0.9988 | 95.0 |
| 1131 | Male | Ts | CB | *Rn18s* | -3.427 | 9.78 | 0.9998 | 95.8 |
| **Mean** |  |  |  |  | -3.407 | 10.00 | 0.9995 | 96.6 |
| **SEM** |  |  |  |  | 0.016 | 0.15 | 0.0001 | 0.6 |

| Supplemental Table 3: Locomotor Activity Test on Postnatal Day 10 (Mean ± SEM) | | | | | |
| --- | --- | --- | --- | --- | --- |
|  | | **Eu,*Dyrk1a*^+^*^/^*^Dox-Cre^** | **Eu,*Dyrk1a*^+^*^/^*^+^** | **Ts,*Dyrk1a*^+/+^*^/^*^+^** | **Ts,*Dyrk1a*^+/+^*^/^*^Dox-Cre^** |
| **Females** | N | 15 | 36 | 12 | 7 |
|  | Total Distance (cm) | 136.5 ± 26.2 | 147.8 ± 21.7**^***^** | 45.2 ± 18.2 | 40.1 ± 10.5 |
|  | % Time  in Center | 28.9 ± 7.1 | 23.1 ± 5.0**^###^** | 67.1 ± 10.8 | 54.6 ± 12.4 |
| **Males** | N | 12 | 28 | 18 | 14 |
|  | Total Distance (cm) | 51.9 ± 13.5 | 97.1 ± 17.4 | 61.9 ± 15.3 | 58.3 ± 16.8 |
|  | % Time  in Center | 28.9 ± 10.7 | 24.2 ± 5.7**^#^** | 45.2 ± 9.0 | 62.5 ± 11.1 |
| *******p<.001, Eu,*Dyrk1a*^+^*^/^*^+^ > Ts,*Dyrk1a*^+^*^/^*^+/+^, Total Distance  **^#^**p<.05, **^###^**p<.001, Eu,*Dyrk1a*^+^*^/^*^+^ < Ts,*Dyrk1a*^+^*^/^*^+/+^, % Time in Center | | | | | |

| Supplemental Table 4: Traversal Benchmarks in Homing Test on Postnatal Day 11 (Number of Pups) | | | | | | |
| --- | --- | --- | --- | --- | --- | --- |
|  | | | Eu,*Dyrk1a*^+^*^/^*^Dox-Cre^ | Eu,*Dyrk1a*^+^*^/^*^+^ | Ts,*Dyrk1a*^+/+^*^/^*^+^ | Ts,*Dyrk1a*^+/+/Dox-Cre^ |
| Females | N | | 14 | 33 | 10 | 6 |
|  | Not reach midline | | 6 (43%) | 6 (18%) | 5 (50%) | 5 (83%) |
|  | Cups Reached | Neither  (includes not reach midline) | 8 (57%) | 15 (45%) | 8 (80%) | 6 (100%) |
|  |  | Home only | 3 | 10 | 2 | 0 |
|  |  | Clean only | 1 | 2 | 0 | 0 |
|  |  | Both | 2 | 6 | 0 | 0 |
| Males | N | | 11 | 25 | 13 | 13 |
|  | Not reach midline | | 4 (36%) | 5 (20%) | 7 (54%) | 4 (31%) |
|  | Cups Reached | Neither  (includes not reach midline) | 6 (55%) | 12 (48%) | 12 (92%) | 5 (38%) |
|  |  | Home only | 2 | 7 | 0 | 5 |
|  |  | Clean only | 1 | 2 | 1 | 2 |
|  |  | Both | 2 | 4 | 0 | 1 |

| Supplemental Table 5: Traversal Benchmarks in Homing Test on Postnatal Day 14 (Number of Pups) | | | | | | |
| --- | --- | --- | --- | --- | --- | --- |
|  | | | Eu,*Dyrk1a*^+^*^/^*^Dox-Cre^ | Eu,*Dyrk1a*^+^*^/^*^+^ | Ts,*Dyrk1a*^+/+^*^/^*^+^ | Ts,*Dyrk1a*^+/+/Dox-Cre^ |
| Females | N | | 14 | 30 | 9 | 6 |
|  | Not reach midline | | 3 (21%) | 0 (0%) | 2 (22%) | 3 (50%) |
|  | Cups Reached | Neither | 3 (21%) | 2 (7%) | 3 (33%) | 3 (50%) |
|  |  | Home only | 3 | 5 | 3 | 1 |
|  |  | Clean only | 0 | 1 | 0 | 0 |
|  |  | Both | 8 | 22 | 3 | 2 |
| Males | N | | 10 | 24 | 13 | 13 |
|  | Not reach midline | | 3 (30%) | 3 (12%) | 2 (15%) | 1 (8%) |
|  | Cups Reached | Neither | 3 (30%) | 5 (21%) | 3 (23%) | 4 (31%) |
|  |  | Home only | 1 | 7 | 3 | 3 |
|  |  | Clean only | 0 | 0 | 1 | 2 |
|  |  | Both | 6 | 12 | 6 | 4 |

**[FIGURES]**

**Supplemental Figure 1**

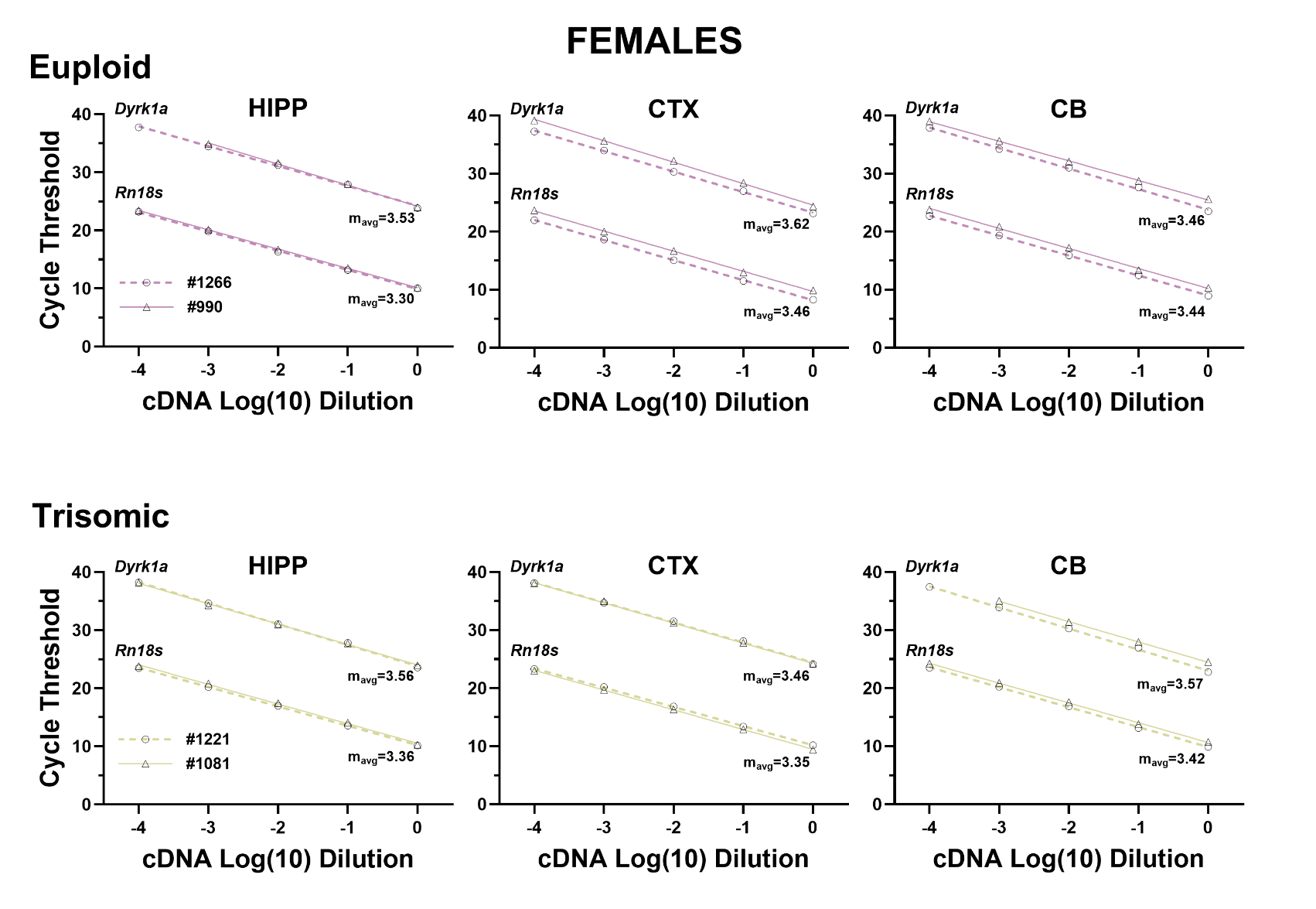

**Supplemental Figure 1: Log-linear serial dilution plots for qPCR amplification of *Dyrk1a* and *Rn18s* of cDNA from P21 hippocampus (HIPP), cerebral cortex (CTX), and cerebellum (CB) of two euploid females and two trisomic females, for validation of qPCR efficiency**. The average slope (m_avg_) is shown for each gene of the two cases for each genotype and brain region. Each point represents the mean of 3 replicates; the average standard deviation of the technical replicates across the five dilutions was 0.136 for *Dyrk1a* and 0.126 for *Rn18s.* The mean (± SEM) efficiency was 92.0% (± 0.9) for *Dyrk1a* and 97.2 (± 0.7) for *Rn18s*. See Supplemental Table 1 for summaries of individual log-linear regression parameters and efficiency calculation for each case and brain region.

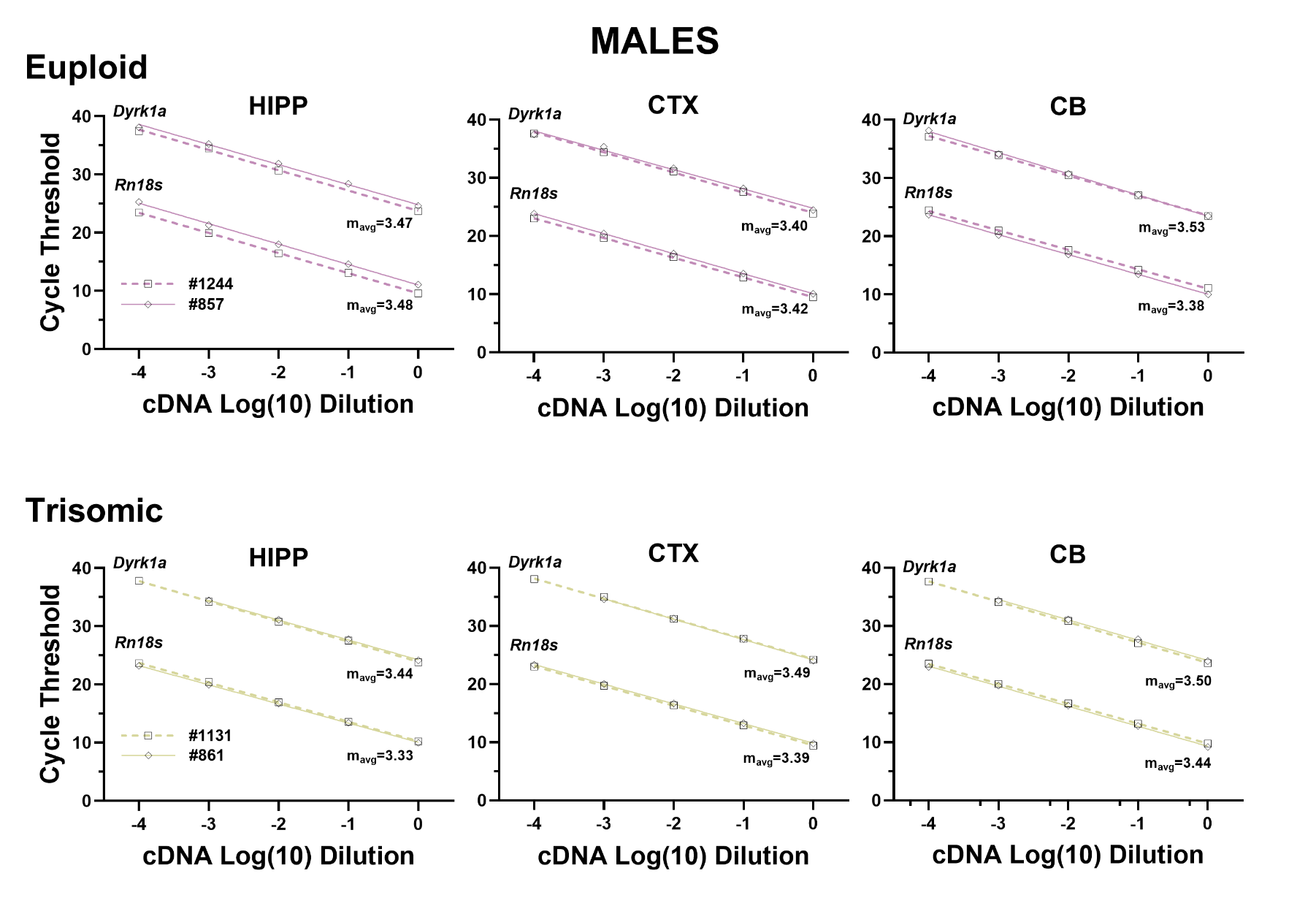
**Supplemental Figure 2**

**Supplemental Figure 2: Log-linear serial dilution plots for qPCR amplification of *Dyrk1a* and *Rn18s* of cDNA from P21 hippocampus (HIPP), cerebral cortex (CTX), and cerebellum (CB) of two euploid males and two trisomic males, for validation of qPCR efficiency.** The average slope (m_avg_) is shown for each gene of the two cases for each genotype and brain region. Each point represents the mean of 3 replicates; the average standard deviation of the technical replicates across the five dilutions was 0.08 for *Dyrk1a* and 0.124 for *Rn18s.* The mean (± SEM) efficiency was 94.2% (± 0.8) for *Dyrk1a* and 96.6 (± 0.67) for *Rn18s*. See Supplemental Table 2 for summaries of individual log-linear regression parameters and efficiency calculation for each case and brain region.

**Supplemental Figure 3**

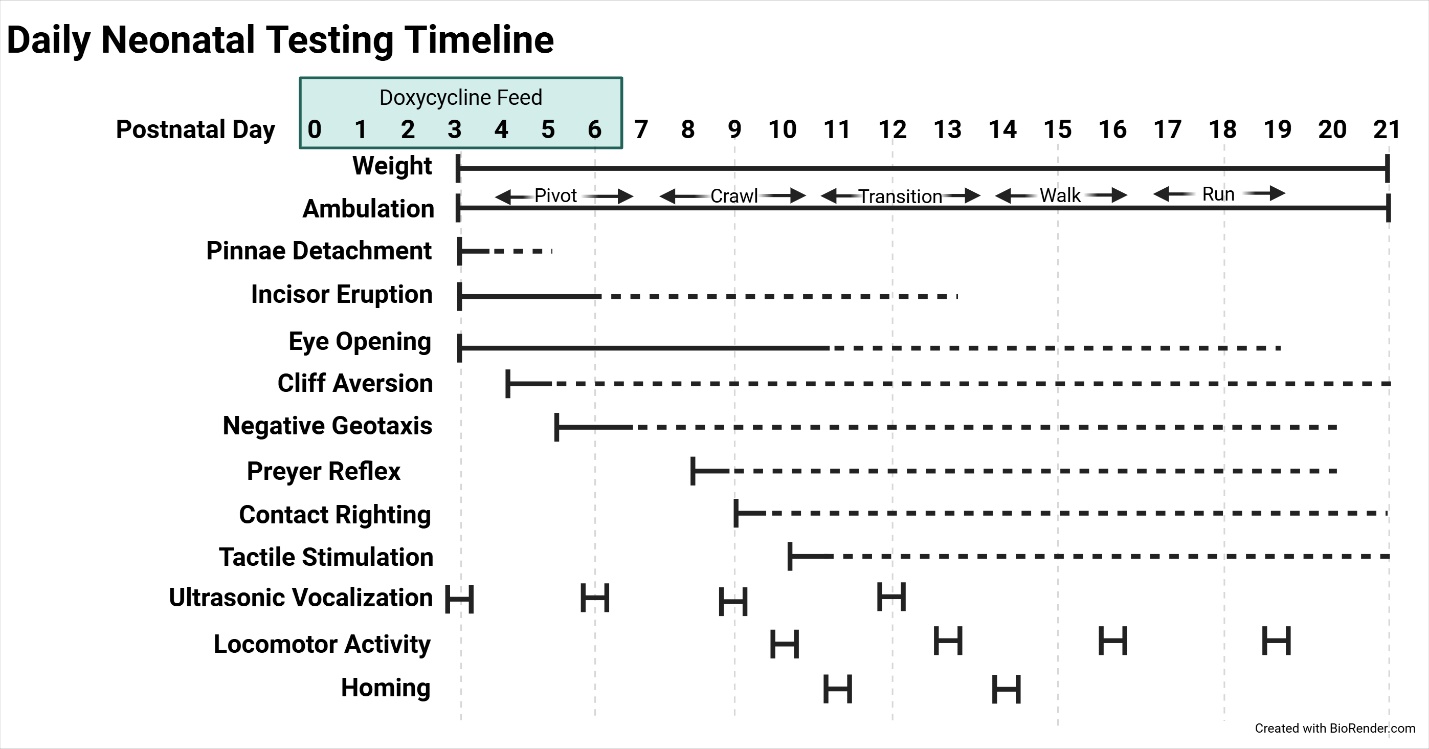

**Supplemental Figure 3: Timeline of Sensorimotor and Neurobehavior Development Assessments.**

**Supplemental Figure 4**

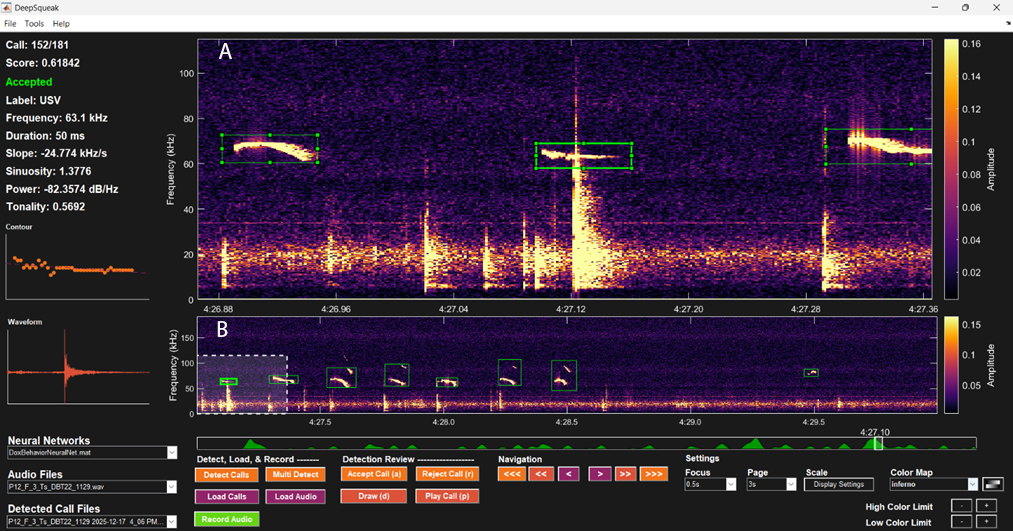

**Supplemental Figure 4: Example of USV sonogram from the DeepSqueak analysis of the audio signals recorded during isolation of a pup.** Analysis of USV sessions used a source file of a neural network that was previously trained by an experienced technician to identify calls and exclude noise. The neural network analysis identified calls emitted during the 5-min session and extracted five parameters of the calls: 1) the number of calls emitted, 2) the average call length (duration), 3) the average peak frequency (kHz) (the maximum intensity frequency), 4) the average principal frequency (kHz) (the most dominant frequency) and 5) the average power (dB/kHz). Note that Panel A of the sonogram shows three calls emitted over ~0.5 secs (from time 4:26.88 to 4:27.36) with the data for the highlighted middle listed to the left. Panel B depicts a longer segment of the sonogram (3 secs, from 4:27.0 to 4:30.0, including the segment in Panel B) showing the 8 identified calls over that time frame. This example 2D graphic representation is from a session of a P12 Ts,*Dyrk1a*^+/+/+^ female mouse.

**Supplemental Figure 5**

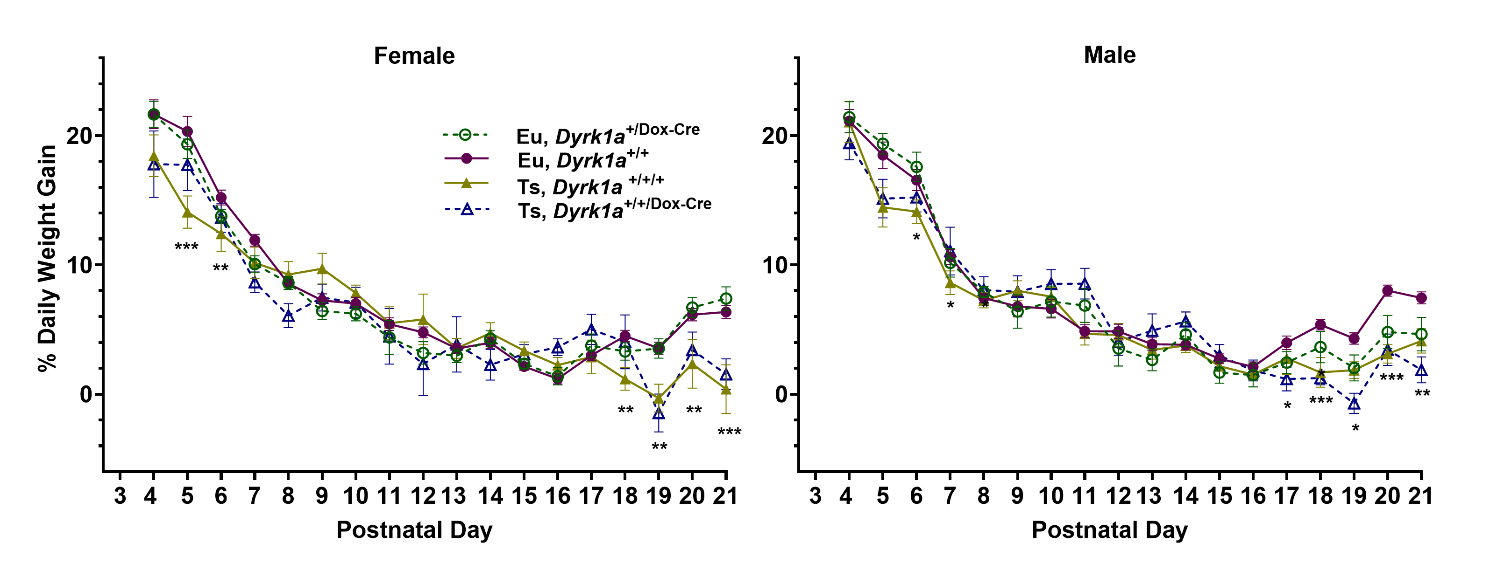

**Supplemental Figure 5.** **Daily Percent Body Weight Change of Female and Male Pups.**

Data presented as mean ± SEM.

*p < 0.05, **p < 0.01, ***p < 0.001, Ts,*Dyrk1a*^+/+/+^ < Eu,*Dyrk1a*^+/+^

**Supplemental Figure 6**

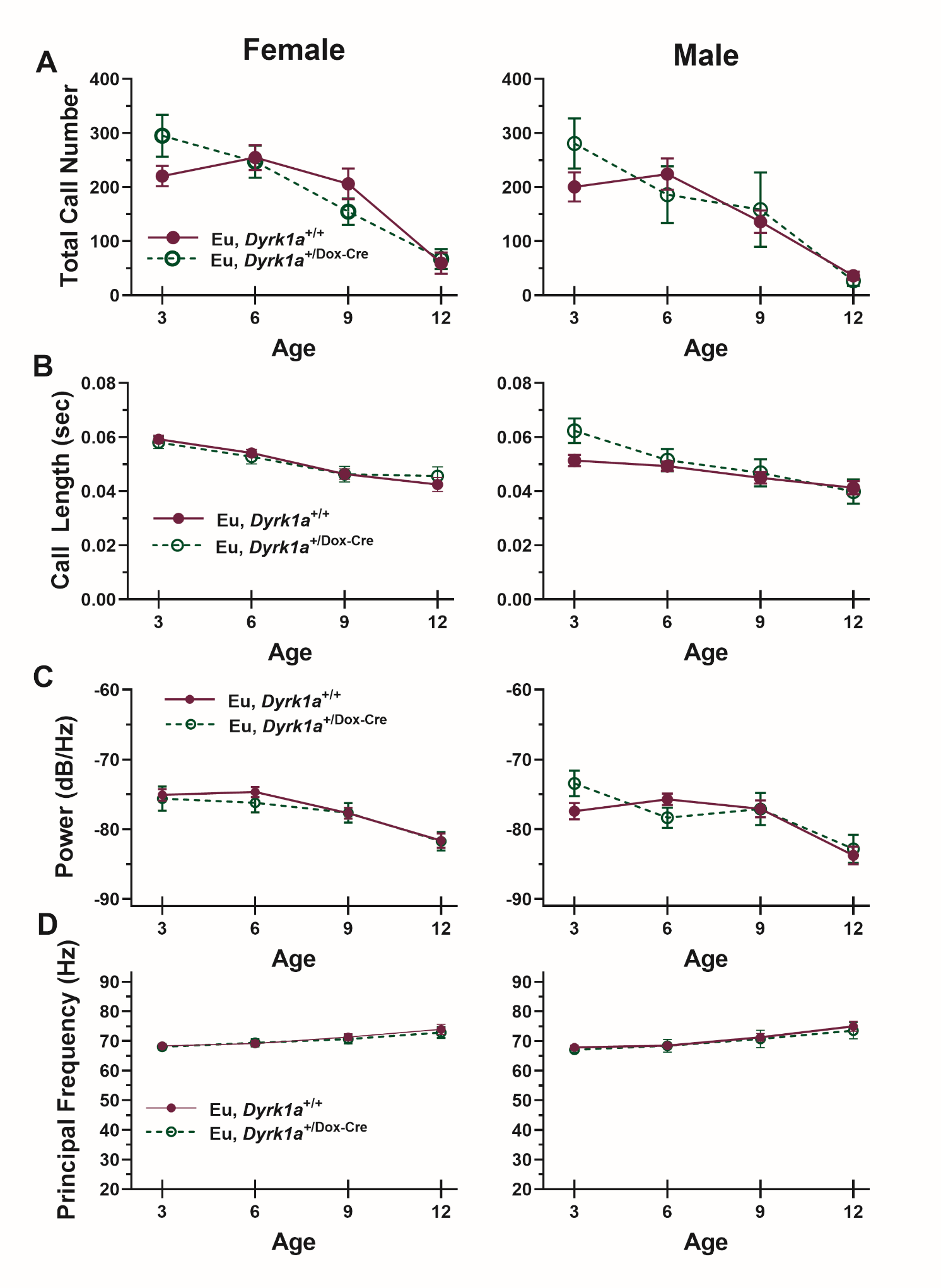

**Supplemental Figure 6: Isolation-induced Ultrasonic Vocalization Measures Comparing Eu,*Dyrk1a*^+/+^ and Eu,*Dyrk1a*^+/Dox-Cre^ Female and Male Mice on Postnatal (P) Days 3, 6, 9, and 12.** There were no significant main or interactive effects of Genotype on any measure.

Data presented as mean ± SEM.

Group numbers: Eu,*Dyrk1a*^+/+^, 36-39 F, 28-30 M; Eu,*Dyrk1a*^+/Dox-Cre^, 14-15 F, 12 M

**Supplemental Figure 7**

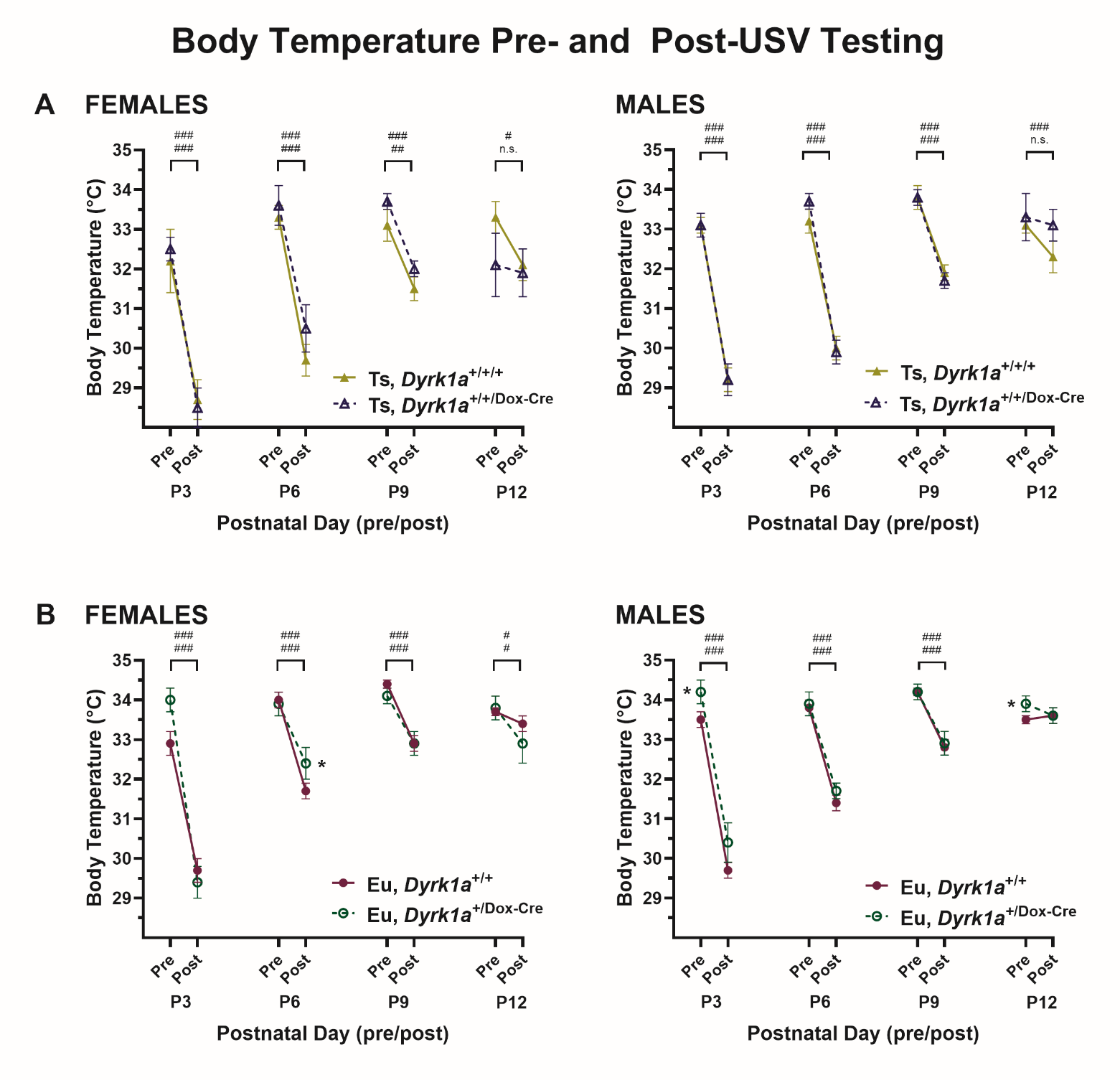

**Supplemental Figure 7. Body Temperatures Before and After Each USV Test Session on P3, P6, P9, and P12 for Female and Male Mice of Ts,*Dyrk1a*^+/+/+^ and Ts,*Dyrk1a*^+/+/Dox-Cre^ Genotypes (A) and of Eu,*Dyrk1a*^+/+^ and Eu,*Dyrk1a*^+/Dox-Cre^ Genotypes (B).** All groups showed significant body temperature loss on P3 that became progressively less severe with increasing age [main effects of day and pre/post and day× pre/post interactions were significant, p’s<.001]. (A): For the a priori directional hypothesis that Ts,*Dyrk1a*^+/+/Dox-Cre^ < Ts,*Dyrk1a*^+/+/+^, there were no significant main or interactive effects of genotype for either sex. (B): For the a priori hypothesis that Eu,*Dyrk1a*^+/Dox-Cre^ ≠ Eu,*Dyrk1a*^+/+^, for females, there was a significant 3-way interaction due mainly to the higher body temperatures of the Eu,*Dyrk1a*^+/Dox-Cre^ group on the P6 post-test (p=.023) and a trend for higher body temperatures on the P3 the pre-test (p=.064). For males, there was a main effect of genotype due mainly to the higher body temperatures of the Eu,*Dyrk1a*^+/Dox-Cre^ group for the pre-test on P3 (p=.037) and P12 (p=.013) as compared to Eu,*Dyrk1a*^+/+^.

^#^p<.05; ^###^p<.001, Sidek post hoc tests: the Post-USV test was significantly reduced relative to the Pre-USV test for the session; n.s.=not significant (the top and bottom symbols refer to the top and bottom groups listed in the panel legend, respectively)

*p<.05; Sidek post hoc tests, Ts,*Dyrk1a*^+/+/+^ group was significantly lower than comparison group

**Supplemental Figure 8**

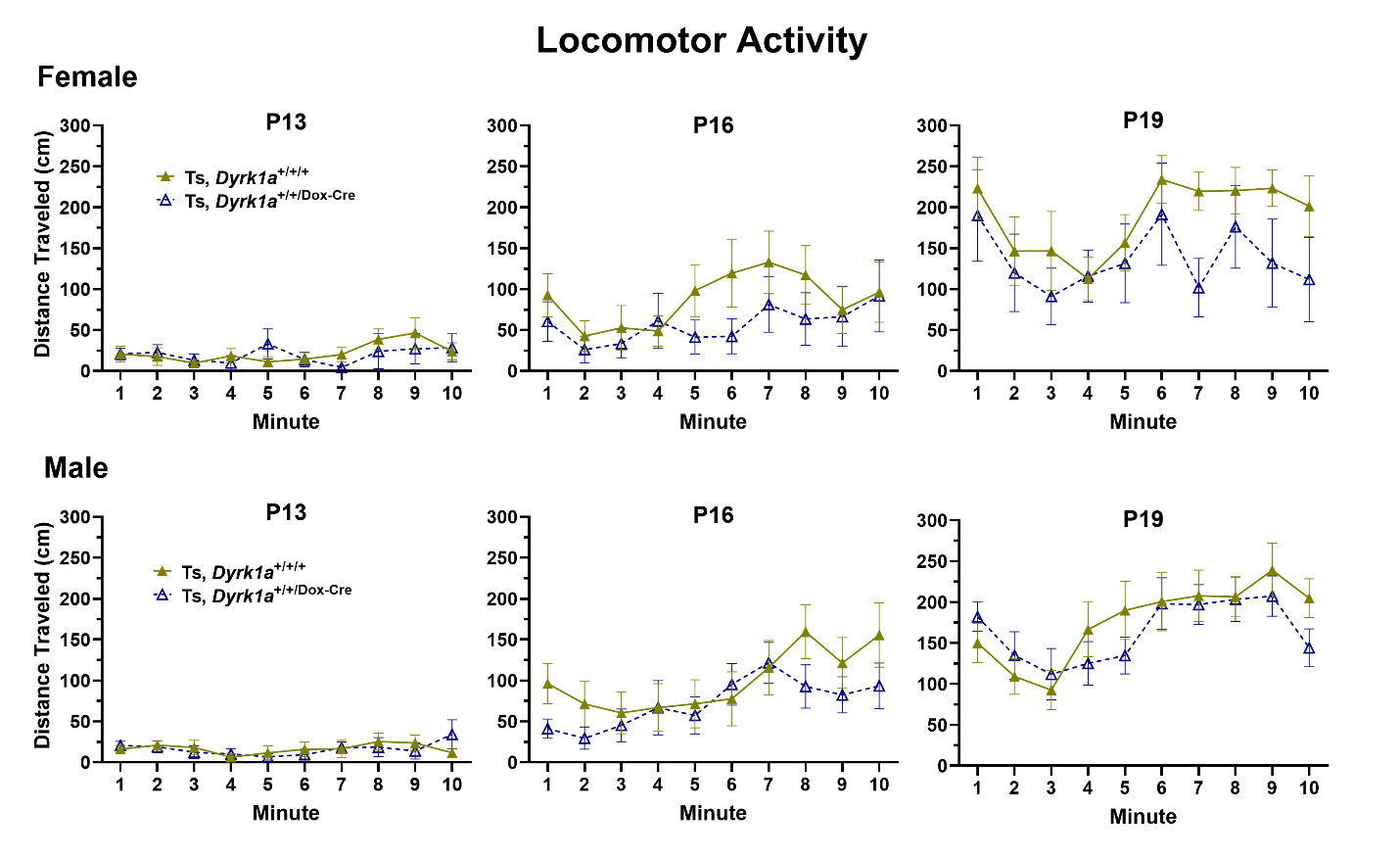

**Supplemental Figure 8: Distance Traveled during the Locomotor Activity Test Comparing Ts,*Dyrk1a*^+/+/+^ and Ts,*Dyrk1a*^+/+/Dox-Cre^ Female and Male mice on Postnatal (P) Days 13, 16, and 19.** Locomotor activity increased across age, but there were no main or interactive effects of genotype for Ts,*Dyrk1a*^+/+/+^ and Ts,*Dyrk1a*^+/+/^**^Dox-Cre^** mice at any age for either sex

Data presented as mean ± SEM.

Group numbers: Ts,*Dyrk1a*^+/+/+^ 11-12 F; 17-19 M; Ts,*Dyrk1a*^+/+/^ **^Dox-Cre^** 8 F; 14 M

**Supplemental Figure 9**

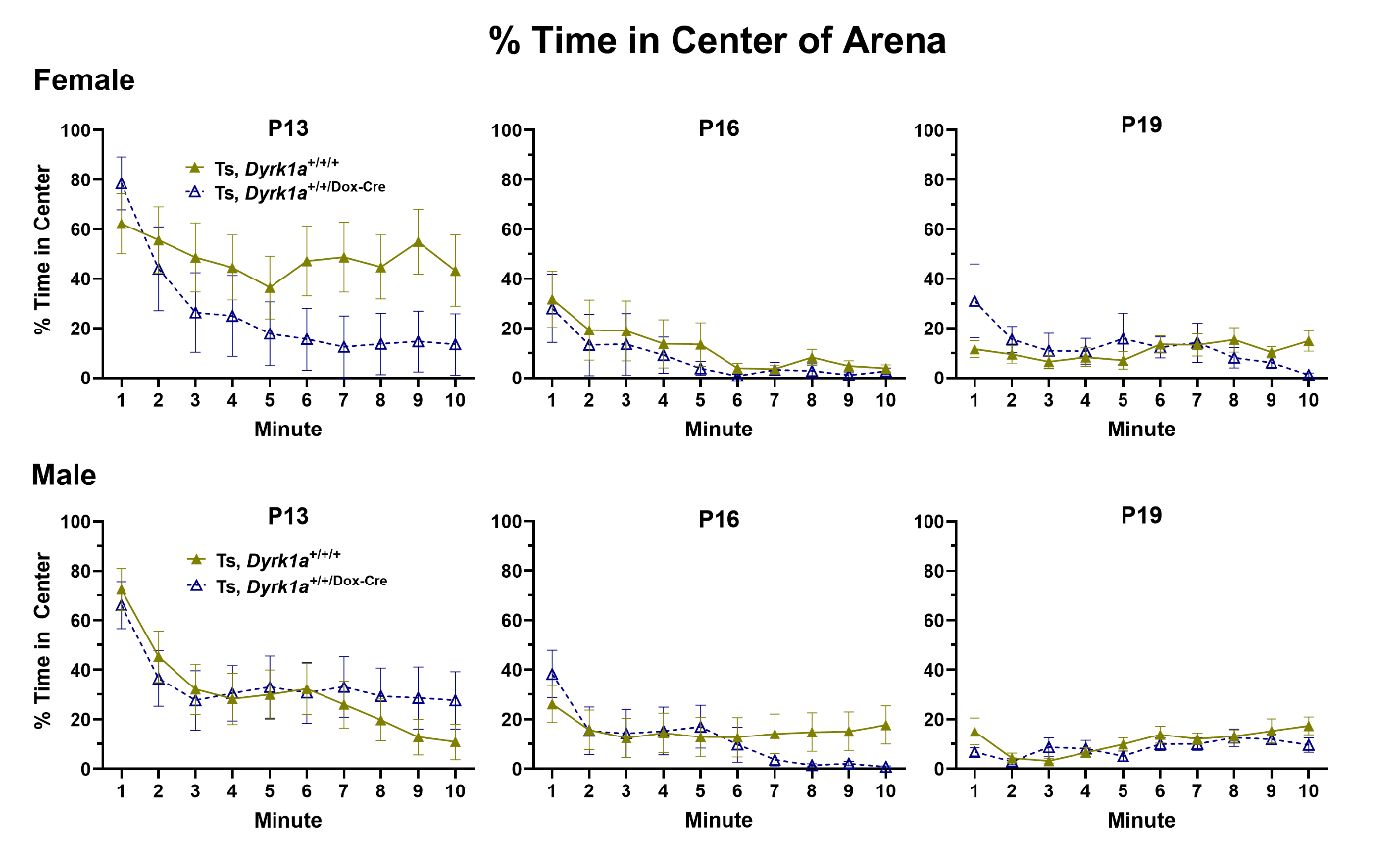

**Supplemental Figure 9: Percent Time in the Center during the Locomotor Activity Test Comparing Ts,*Dyrk1a*^+/+/+^ and Ts,*Dyrk1a*^+/+/Dox-Cre^ Female and Male Mice on Postnatal (P) Days 13, 16, and 19.** On P13, the differences in time in center between Ts,*Dyrk1a*^+/+/+^ and Ts,*Dyrk1a*^+/+/^ ^Dox-Cre^ mice did not reach significance for either sex (the trend for a Genotype X Minute interaction in females was not significant, p=.063). On P16, time in center declined after the first minute in all groups. There were no significant main or interactive effects of genotype in females. but the greater reductions in male Ts,*Dyrk1a*^+/+/^**^Dox-Cre^** mice as compared to Ts,*Dyrk1a*^+/+/+^ mice in the last four minutes yielded a Genotype X Minute interaction (p=.033); nevertheless, the individual comparisons on minutes 7-10 did not reach significance. On P19, males and females of both groups largely avoided the center zone after the first minute mice, with average time in center all <20% on the remaining 9 minutes. There were no significant main or interactive effects of genotype on P19.

Data presented as mean ± SEM.

Group numbers: Ts,*Dyrk1a*^+/+/+^ 11-12 F; 17-19 M; Ts,*Dyrk1a*^+/+/^**^Dox-Cre^** 8 F; 14 M

**Supplemental Figure 10**

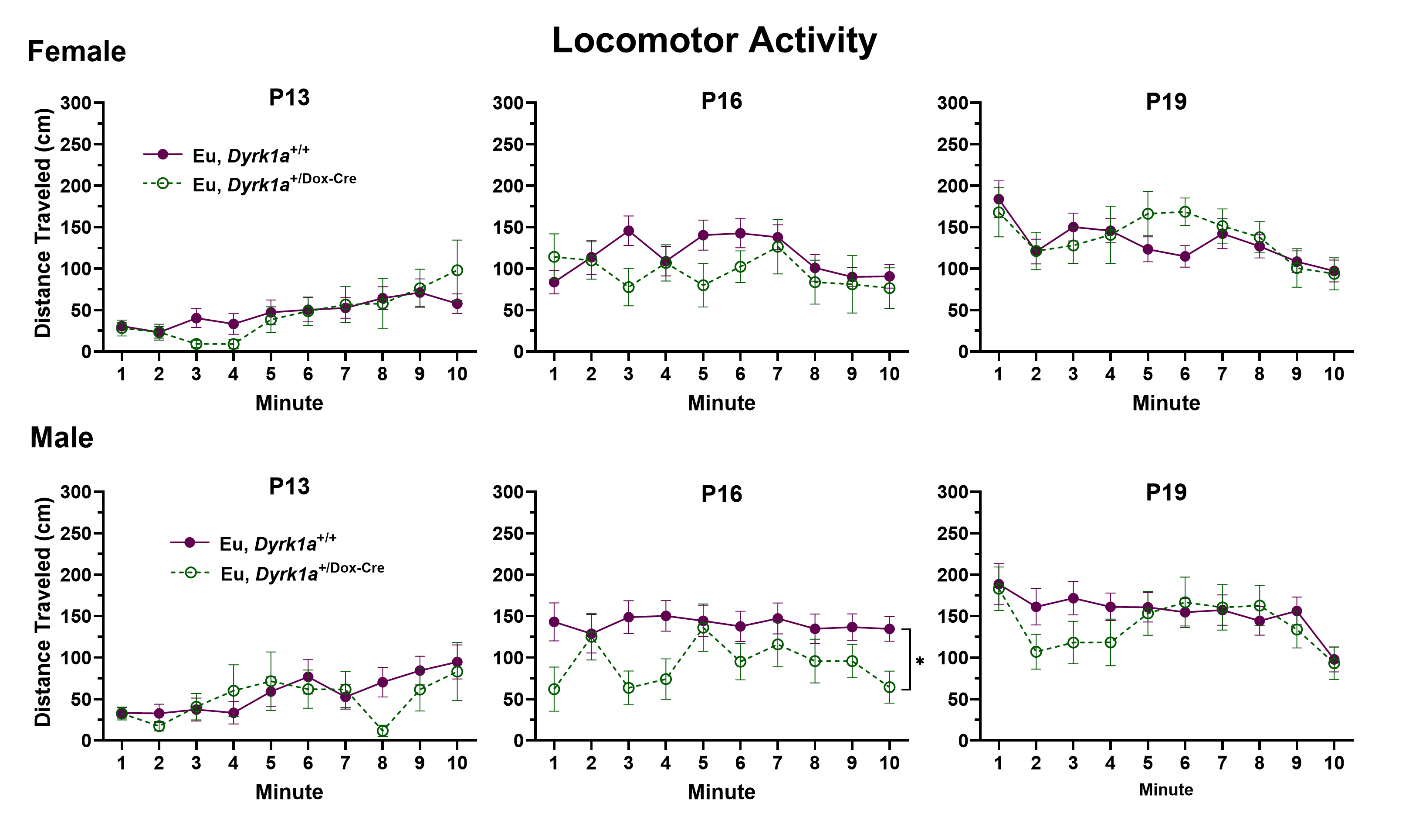

**Supplemental Figure 10: Distance Traveled during the Locomotor Activity Test Comparing Eu,*Dyrk1a*^+/+^ and Eu,*Dyrk1a*^+/Dox-Cre^ Female and Male Mice on Postnatal (P) Days 13, 16, and 19.** Locomotor activity increased across age, but there were no significant differences between Eu,*Dyrk1a*^+/+/+^ and Eu,*Dyrk1a*^+/+/^ **^Dox-Cre^** mice at any age for either sex.

Data presented as mean ± SEM.

Group numbers: Eu,*Dyrk1a*^+/+/+^ 36-37 F, 31 M; Eu,*Dyrk1a*^+/+/^ **^Dox-Cre^** 12-15 F, 12 M

**Supplemental Figure 11**

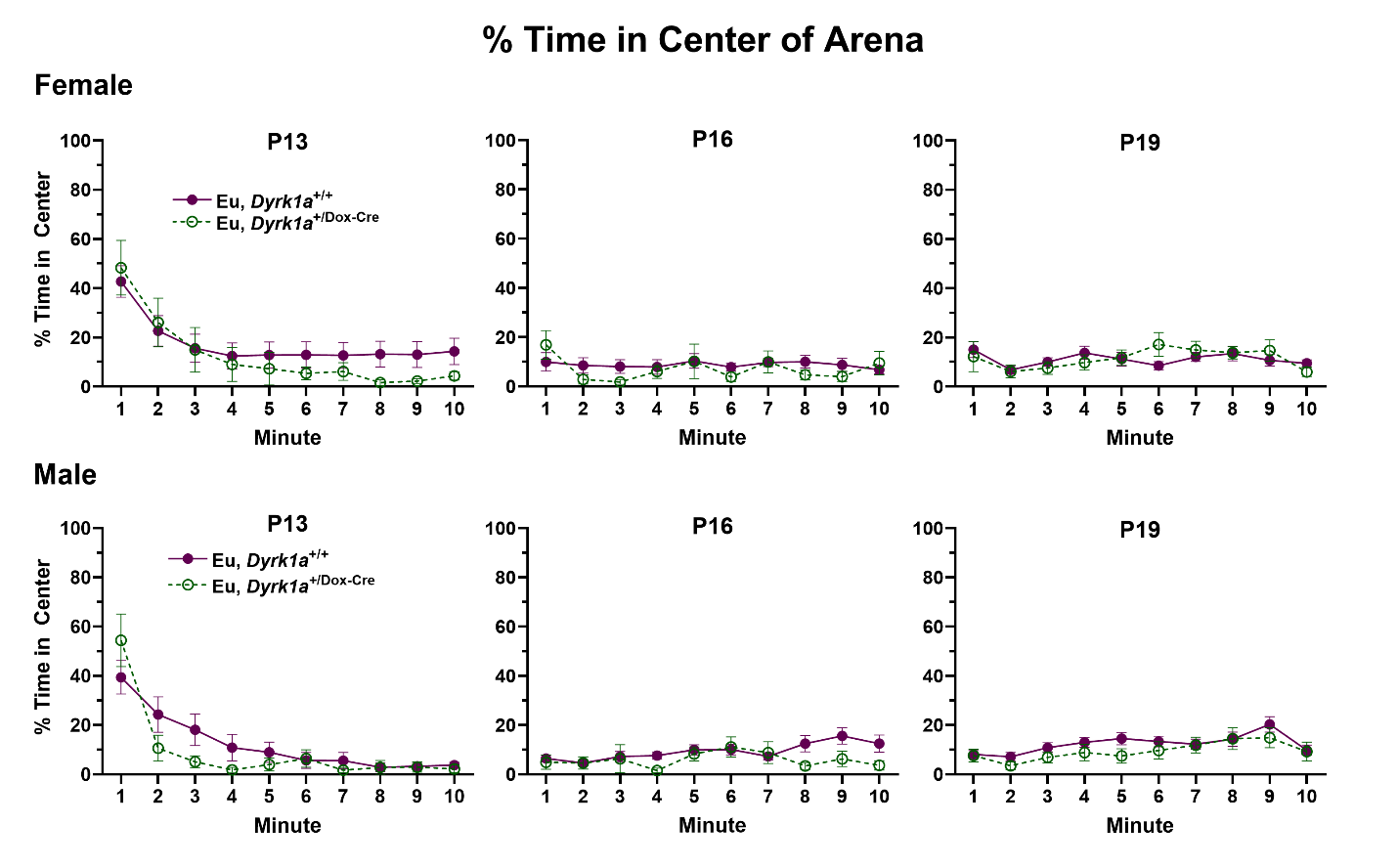

**Supplemental Figure 11: Percent Time Spent in Center during the Locomotor Activity Test Comparing Eu,*Dyrk1a*^+/+^ and Eu,*Dyrk1a*^+/ Dox-Cre^ Female and Male Mice on Postnatal (P) Days 13, 16, and 19.** Time in center declined over the P13 session and for P16 and P19 never exceeded 20% on average for any group for any minute. There were no significant differences between Eu,*Dyrk1a*^+/+/+^ and Eu,*Dyrk1a*^+/+/^ **^Dox-Cre^** mice at any age for either sex.

Data presented as mean ± SEM.

Group numbers: Eu,*Dyrk1a*^+/+/+^, 36-37 F; 31 M; Eu,*Dyrk1a*^+/+/^ **^Dox-Cre^**, 12-15 F; 12 M
